# Cellular-Resolution Spatial Transcriptomics Reveals Laminar VSMC Phenotypic Remodeling and a Hypoxic Medial Core in Human Thoracic Aortic Dissection

**DOI:** 10.64898/2026.07.13.738344

**Authors:** Mary A. Siki, Krishna Gajera, Patryk A. Dabek, M. Grady Freeman, Yuling Zhu, Pamela K. Woodard, Alexander A. Brescia, Benjamin D. Humphreys, Katherine M. Holzem

## Abstract

**Objectives:** Acute aortic syndromes (AAS) carry high short-term mortality and long-term morbidity. Hypertension is the predominant risk factor, yet no established tools identify patients at imminent risk. Despite known vascular smooth muscle cell (VSMC) phenotype switching with hypertension, the medial role in sporadic AAS is given comparatively less emphasis, compared to heritable aortopathies. We sought to define the VSMC landscape in human AAS and identify spatial remodeling programs associated with interlamellar separation.

**Methods:** We collected human ascending aorta specimens from 8 patients undergoing open surgical AAS repair and 5 donors after brain death. Using Xenium in situ spatial transcriptomic profiling, we generated cellular-resolution maps of human ascending aorta. Tissue sections were subsequently stained for histology and immunohistochemistry.

**Results:** In AAS, we identified extensive VSMC remodeling organized into distinct laminar domains across the aortic media, with unique VSMC states supported by gene-expression module scoring. Across patients, we identified a reproducible mid-medial core of hypoxia-adapted VSMCs spatially distinct from acute injury-associated states bounding the false lumen which were characterized by carbonic anhydrase 9 immunoreactivity and reduced α-smooth muscle actin expression. Comparatively, non-dissected donor ascending aortas contained fewer remodeled VSMC states, minimal transmural striation, and no discrete regions of hypoxia-adapted VSMCs.

**Conclusions:** These findings define a spatially organized medial remodeling landscape in AAS and identify hypoxia-associated VSMC modulation as a potential feature of medial domains associated with heightened vulnerability to interlamellar separation. Defining these remodeling domains may inform improved risk stratification and experimental models that better recapitulate human AAS pathology.

**Ultra-mini Abstract:** We sought to define the vascular smooth muscle cell (VSMC) landscape in human acute aortic syndrome (AAS) using spatial transcriptomics. AAS demonstrated extensive laminar VSMC remodeling, including a mid-medial core of hypoxia-adapted VSMCs. These findings identify hypoxia-associated VSMC modulation as a potential feature of medial domains vulnerable to interlamellar separation.

AAS demonstrated extensive laminar VSMC remodeling, including a mid-medial core of hypoxia-adaptation hypoxia-associated VSMC modulation as a potential feature of medial domains vulnerable to interlamellar separation.

## Introduction

Acute aortic syndromes (AAS) involving the ascending aorta, including acute type A ascending aortic dissection (ATAAD) and acute type A intramural hematoma (ATAIMH), can be acutely fatal vascular emergencies with poor long-term prognoses. In-hospital mortality approaches 25% for both conditions despite emergency repair, and ten-year survival after acute event approximates 60%.^1–5^ Even for survivors, lifelong surveillance is required, as aortic reintervention reaches 11.6% at 5 years and 16.0% at 10 years, and a persistent false lumen quadruples the risk of distal aortic events.^6–8^ Hypertension (HTN), present in 70% to 85% of affected patients, is the predominant modifiable risk factor but has limited ability to discriminate individual risk.^5,9^ Importantly, no early diagnostics exist for prophylactic intervention, and the majority of research to date has focused on medical and surgical management after irreversible AAS has occurred.

ATAAD is conventionally attributed to an intimal tear that extends through the internal elastic lamina, allowing pressurized blood to enter the aortic media and propagate a false lumen.^10^ Although this model explains false lumen formation after an intimal breach, it does not fully explain why dissection occurs preferentially in heritable aortopathy nor does it reconcile the similar pathology of ATAIMH. In either case of sporadic AAS, medial susceptibility remains underemphasized, despite the established role of HTN in driving adverse aortic remodeling and vascular smooth muscle cell (VSMC) phenotypic modulation.^11,12^ Here, we examine sporadic AAS as a disease of the aortic media, in which maladaptive VSMC states may define the tissue context in which pathological medial separation occurs and provide the underlying pathologic substrate in ATAAD and ATAIMH.

We hypothesized that spatially distinct remodeled domains within the aortic media are associated with AAS. The cellular events that distinguish specific regions of the dissected media are poorly defined, but regional differences in wall compliance and failure strength are known to influence where interlamellar tearing begins. ^13–15^ Hypoxia has also been proposed as a feature of the remodeled medial environment associated with dissection risk.^16^ Together, these observations support a model in which regional microenvironmental and biomechanical heterogeneity may define the medial substrate for interlamellar separation. Although dissociated single-cell (sc) and single-nucleus (sn) approaches have identified distinct cellular states in aortic disease, spatial analyses of human AAS have remained limited in cellular resolution and preservation of tissue architecture.^17^ We therefore used cellular-resolution spatial transcriptomics to define how VSMC phenotypic states and remodeling programs are organized across the human AAS wall and generated a cellular-resolution spatial map of pathologic and donor human ascending aorta and define VSMC states and tissue domains associated with medial remodeling in AAS.

## Methods

### Human subject identification and recruitment

The study was evaluated and approved by the Washington University Institutional Review Board (IRB #202603063). Adult AAS patients undergoing open surgical repair at Barnes-Jewish Hospital, St. Louis, MO were consented for enrollment in the institutional Vascular Surgery Biobank (IRB #201309043). Non-identifying clinical information was collected via electronic medical record review per biobank protocol. Patients with history of prior cardio-aortic surgery were excluded, as iatrogenic causes for ascending aortic pathology could not be ruled out. Additionally, donor control (DC) aortic specimens were collected from organ donors who consented to research tissue collection through partnership with Mid-America Transplant Services (St. Louis, MO).

### Tissue collection and Xenium processing

Ascending aorta segments were immediately recovered in the operating room by a member of the study team after excision for AAS specimens or after perfusate administration for DC specimens. When able, anatomic orientation (i.e. proximal edge, greater curve) was noted at the time of recovery. For analysis, we selected tissue segments spanning the margin of dissection or interlamellar separation together with the adjacent intact, full-thickness aortic wall, when present. Because these paired regions most commonly localized to the lesser curvature, the resulting dataset preferentially represents the lesser rather than the greater curvature. Given this localization in AAS specimens, DC aortic tissue regions were also processed from the lesser curvature for more direct comparison. Tissues were gently rinsed with saline, without disrupting any attached hematoma if present, fixed in 10% formalin solution for 18-24 hours, and then transferred to 70% EtOH solution for storage. Next, specimens were paraffin embedded with orientation documented, cut into 6 μm transverse sections, and carefully mounted onto Xenium slides.

A custom 480-gene Xenium panel was designed for analysis. The panel consisted of genes specific for vascular biology and pathophysiology including distinct inflammatory, hypoxia, extracellular matrix remodeling, osteogenesis, and cell signaling markers (Supplemental Table 1). Xenium slides were deparaffinized and decrosslinked followed by probe hybridization and rolling amplification. Slides were then stained using the Cell Segmentation Kit (10x Genomics, Pleasanton, CA, USA), which uses a variety of protein markers to help define cell boundaries. Slides were analyzed and imaged with the Xenium analyzer, and data were visualized with Xenium Explorer software. Sequencing was performed using the Illumina NovaSeq X Plus at the McDonnell Genome Institute at WashU.

### Histology and Immunohistochemistry

After Xenium processing, sections were stained with hematoxylin and eosin (H&E) according to 10x Genomics protocols.^18^ Neighboring 6μm specimens were sectioned onto charged histology slides for immunohistochemical (IHC) staining. After deparaffinization and antigen unmasking, we incubated the tissue sections in mouse anti-smooth muscle actin alpha (SMA-α) primary antibody (clone CGA7, Millipore Sigma, Cat# A7607) at a 1:200 dilution and rabbit anti-carbonic anhydrase 9 (CA9) primary antibody (clone EPR23055-5, abcam, Cat# AB243660-1007) at a 1:500 dilution with 3% Bovine Serum Albumin (BSA)-Tris-Buffered Saline with Tween 20 (TBST) blocking buffer at 4° Celsius (C) overnight. These were subsequently stained with donkey anti-mouse IgG (H+L), Alexa Fluor 546 conjugate (Thermo Fisher Scientific, Cat# A10036, RRID: AB_11180613) at a 1:200 dilution and donkey anti-rabbit IgG (H+L), Alexa Fluor 488 conjugate (Thermo Fisher Scientific, Cat# A32790, RRID: AB_2762833) at a 1:500 dilution with 3% BSA-TBST blocking buffer at room temperature for 90 min. The tissue sections were treated with TrueView Autofluorescence Quenching Kit (Vector, Newark, CA, USA) followed by VECTASHIELD Vibrance® Antifade Mounting Medium with 4′,6-diamidino-2-phenylindole (DAPI). Slides were visualized and images obtained by confocal microscopy using Nikon C2+ Eclipse (Fig. 2b) and Nikon ECLIPSE Ti2.

### Spatial transcriptomic processing and clustering

Multiple processing pipelines were evaluated to identify a robust cellular spatial computational pipeline (Supplemental Methods; parameters in Supplemental Table 2). Cell segmentation was performed with Baysor using nucleus boundaries from the Xenium default segmentation, and cell x gene count matrices were constructed as AnnData objects in Scanpy. After quality-control filtering, expression was normalized and log_1_p-transformed. Batch correction across tissue sections was applied using Harmony on the principal component embedding.^19^ Integration quality was validated using batch silhouette score, integration, Local Inverse Simpson’s Index, and clustering (Supplemental Methods). Spatially-aware dimensionality reduction was then performed with the Building Aggregates with a Neighborhood Kernel and Spatial Yardstick (BANKSY) method, which augments each cell’s expression profile with a weighted average of its spatial neighbors to incorporate tissue context.^20^

Leiden clustering was performed on the Harmony-corrected BANKSY embedding (λ=0.2), and Uniform Manifold Projection (UMAP) was used for visualization across all cells (AAS = 877,085 cells, DC = 237,959 cells). AAD and DC datasets were Harmony integrated separately, and Leiden resolutions were tuned to each individual dataset. Because these datasets comprised biologically distinct groups, this ensured that subtle cell states unique to either group were retained rather than artificially homogenized across the disease–control divide. Leiden resolutions were chosen to best resolve VSMC phenotypic diversity within each while avoiding over-clustering (AAS: 1.2, DC: 0.7); therefore, in some cases, rare immune cell populations could not be fully resolved at the same resolution. In addition to cellular-level clustering, mural spatial domains were identified in both AAS and DC samples using increased lambda weighting (λ=0.8).

For each gene, the average log_2_ fold change in expression, together with the percentages of cells expressing the gene within and outside a given cluster, were used to identify marker genes, assign cluster cell types, and infer activated or repressed cellular programs at the time of tissue fixation. Cluster identities were independently reviewed by two investigators, with large language models used as an adjunct aid. Final labels were determined by human consensus.

### VSMC Phenotype Depth Relationship Analysis

In order to determine whether VSMC phenotypes occupy different spatial positions across the aortic wall, a computational processing pipeline was created that leveraged a Laplace-equation solver to identify individual cellular coordinates along a gradient from lumen to adventitia (for DCs) or dissection plane (for AASs). Spatial boundaries were initially hand-traced in QuPath on DAPI morphology image which was then snapped to the segmented-cell cloud.^21^ Lumen boundary was assigned a depth of 0 and adventitia or dissection plane was assigned a depth of 1. VSMC clusters were then masked onto the depth analysis. Median depth of each phenotype was determined. Pooled analysis of cells were used to create mountain plots.

## Results

### Spatial transcriptomic profiling robustly resolves aortic cell types

Computational analysis of 15 aortic sections, sampled from the dissection margin and intact adjacent wall when present, from 8 unique AAS patients (Table 1) revealed 28 cell type clusters. There were 3 clusters with < 2,000 cells each, which represented rare artificial expression profile mixtures and were excluded from our primary analyses. UMAP including these rare clusters are shown in Supplemental Figure 1. From the 25 remaining clusters, we identified 14 clusters of presumptive VSMC lineage, with the remaining clusters comprised of 6 inflammatory cell, 1 fibroblast, 1 pericyte, 1 ECs, 1 adipocyte, and 1 Schwann cell types. Spatial maps of a representative diseased aortic segment, oriented from the adventitia to the lumen, show all cell types, all VSMCs, and all non-VSMCs (Fig. 1A, i–iii). For each spatial plot, the boxed area is displayed at 4× magnification to provide higher-resolution visualization of cellular organization. These maps reveal a striking transmural, striated distribution of VSMC subtypes across the media of the dissected aorta, whereas most non-VSMCs localize to the adventitia, border the dissection flap, or line the luminal surface. UMAP (Fig. 1B) demonstrates a bilobed spatial relationship with mesenchymal lineage clusters toward the right-hand side of the plot and other clusters toward the left. In Fig. 1C, dot plot of functional gene module scores is displayed, highlighting the diverse VSMC transcriptional modulation, with clusters 3 and 9 demonstrating activation of hypoxia-adapted programs.

**Figure 1.**
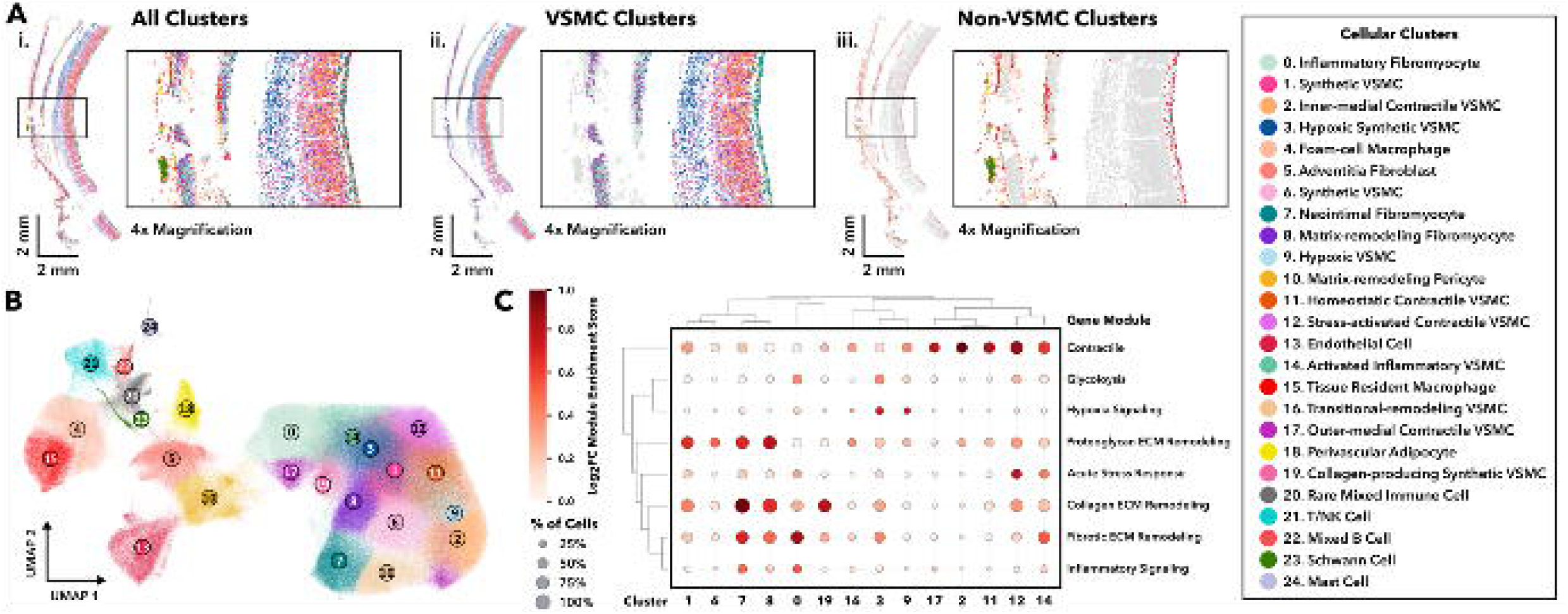
Harmony-integrated spatial transcriptomic analysis of 15 aortic tissue regions from 8 unique AAS patients. **(A)** Faceted spatial plots of a representative AAS segment with **(i)** all clusters, **(ii)** Vascular Smooth Muscle Cell (VSMC) clusters, **(iii)** non-VSMC clusters. Spatial plots displayed adventitia to lumen from left to right. Boxes highlight 4x magnified regions displayed next to each spatial plot. **(B)** Uniform Manifold Projection (UMAP) showing 25 unique cellular clusters. **(C)** Gene module score dot plot of VSMC types demonstrating distinct phenotypic modulation among clusters.

**Table 1:** Acute aortic syndrome (AAS) patient characteristics.

| Variable | AAS_1 | AAS_2 | AAS_3 | AAS_4 | AAS_5 | AAS_6 | AAS_7 | AAS_8 |
| --- | --- | --- | --- | --- | --- | --- | --- | --- |
| Age | 63 | 65 | 60 | 71 | 80 | 58 | 70 | 70 |
| Sex | M | F | F | F | F | F | F | M |
| Hypertension | Yes | Yes | Yes | No | Yes | Yes | Yes | No |
| Hyperlipidemia | Yes | No | Yes | No | Yes | No | Yes | No |
| Coronary Artery Disease | Yes | Yes | Yes | No | Yes | No | Yes | Yes |
| Diabetes | No | Yes | Yes | No | No | No | Yes | No |
| Chronic Kidney Disease | No | No | No | No | No | No | No | No |
| COPD | Yes | Yes | Yes | No | Yes | No | Yes | No |
| Smoking Status | Former | Former | Current | Never | Former | Current | Former | Never |
| Dissection Type (DeBakey) | I | II | II | I | II (IMH) | II | I | I |
| Time from Symptom Onset to Surgery | <24h | 7 d | <24h | <24h | <24h | <24h | <24h | <24h |
| Ascending Aorta diameter at time of dissection (cm) | 5.0 | 4.6 | 4.8 | 5.6 | 5.9 | 4.0 | 4.7 | 4.8 |
| Zone of Tear | 0 | 0 | 0 | 1 | N/A (IMH) | 0 | 0 | 0 |
| False Lumen Location | GC | GC | GC | GC | N/A (IMH) | GC | GC | GC |
| Zone of Distal Extent | 10 | 1 | 0 | 11 | 0 | 0 | 9 | 11 |
| Mortality | Yes | No | No | No | No | No | Yes | No |

### Donor aortas lack major laminar remodeling and hypoxia-adapted transcriptional VSMC programs

We also performed spatial transcriptomic profiling on non-dissected DC aortas from 5 unique patients (Table 2). Harmony-integrated analysis of 5 distinct DC lesser-curvature aortic segments from 5 donors revealed 16 unique cell niches. Of these, 6 were distinct VSMC subtypes, with the remaining 10 clusters comprised 4 inflammatory cell types, 2 fibroblast, 2 endothelial, 1 pericyte, and 1 Schwann cell populations. Spatial plots of all cell types (Fig. 2A-i), all VSMCs (Fig. 2A-ii), and all non-VSMCs (Fig. 2A-iii), each with a boxed region shown at 4× magnification, reveals minimal transmural striation of VSMC phenotypes compared with AAS specimens in Fig. 1. Similar to AAS aortas, the UMAP (Fig. 2B) has a bilobed appearance, with mesenchymal lineage clusters toward the right and other cell populations toward the left. Gene-expression module scoring (Fig. 2C) demonstrates upregulation of synthetic programs and acute-response transcription factors in selected VSMC clusters, without enrichment of inflammatory, glycolytic, or hypoxia-response programs.

**Figure 2.**
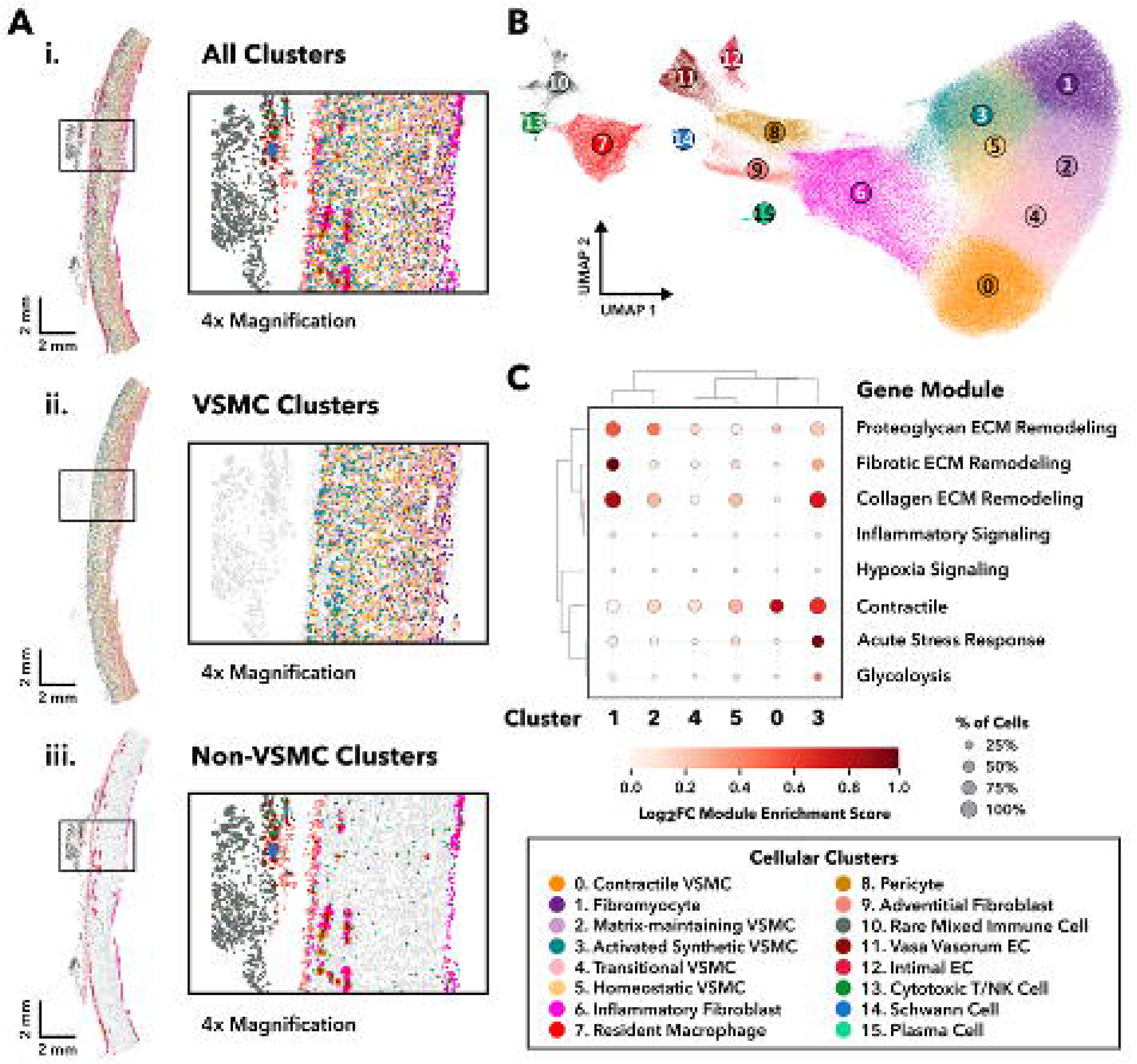
Harmony-integrated spatial transcriptomic analysis of 5 ascending aortic tissue regions from 5 unique DCs. **(A)** Faceted spatial plots of a representative DC segment with **(i)** all clusters, **(ii)** VSMC clusters, **(iii)** non-VSMC clusters. Spatial plots displayed adventitia to lumen from left to right. Boxes highlight 4x magnified regions displayed next to each spatial plot. **(B)** Uniform Manifold Projection (UMAP) showing 16 unique cellular clusters. **(C)** Gene module score dot plot of VSMC types arranged by dendrogram relationship, demonstrating no upregulation of hypoxia-associated or inflammatory signaling programs in DC aortas.

**Table 2:**
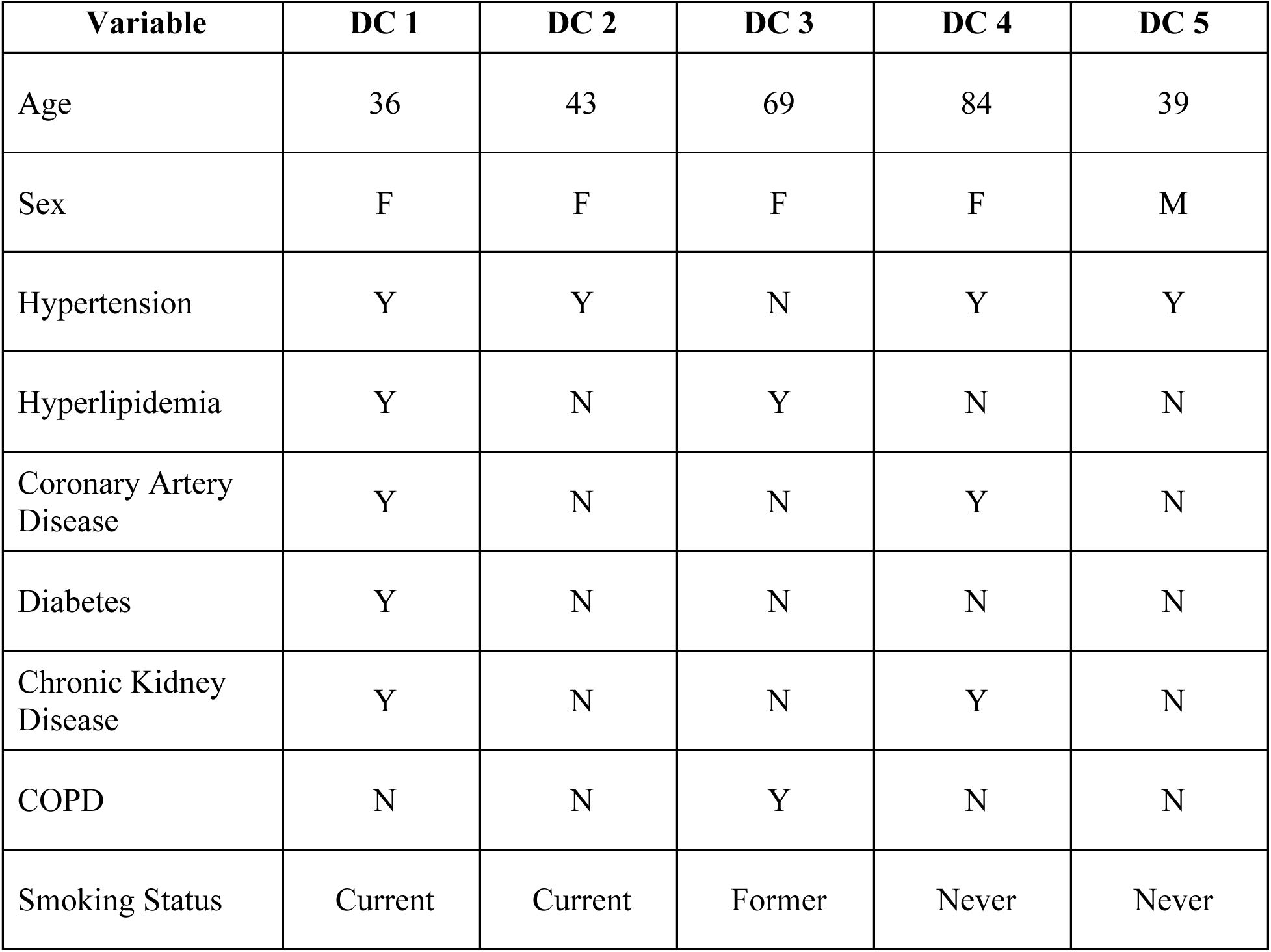
Donor control (DC) patient characteristics.

### IHC validation of prolonged VSMC hypoxia signaling in AAS but not DC tissue

To visualize the tissue architecture and further evaluate hypoxia-adapted tissue regions in AAS compared with DC specimens, H&E and IHC were performed on representative tissue specimens. CA9 was selected as a marker of sustained hypoxia-associated signaling, and co-labeling with SMA-α was used to determine whether hypoxia-associated regions coincided with VSMC phenotypic modulation, as reflected by reduced expression of this key contractile marker Fig. 3A shows representative images from an AAS specimen: (i) H&E staining; (ii) dual immunofluorescence labeling for SMA-α and CA9 with DAPI nuclear counterstaining; (iii) α-SMA-α labeling alone; and (iv) CA9 labeling alone. An inner medial region of CA9 immunoreactivity was seen with associated loss of SMA-α expression consistent with spatial transcriptomic findings from Fig. 1. By contrast, representative H&E and IHC images for a DC aortic specimen are shown in Fig. 3B: (i) H&E staining; (ii) co-labeling for SMA-α and CA9 with DAPI counterstaining; (iii) SMA-α labeling alone; and (iv) CA9 labeling alone. Minimal CA9 labeling is demonstrated within the DC specimen with an associated uniform transmural SMA-α expression, suggesting lack of sustained VSMC hypoxia adaptation and phenotype switching.

**Figure 3.**
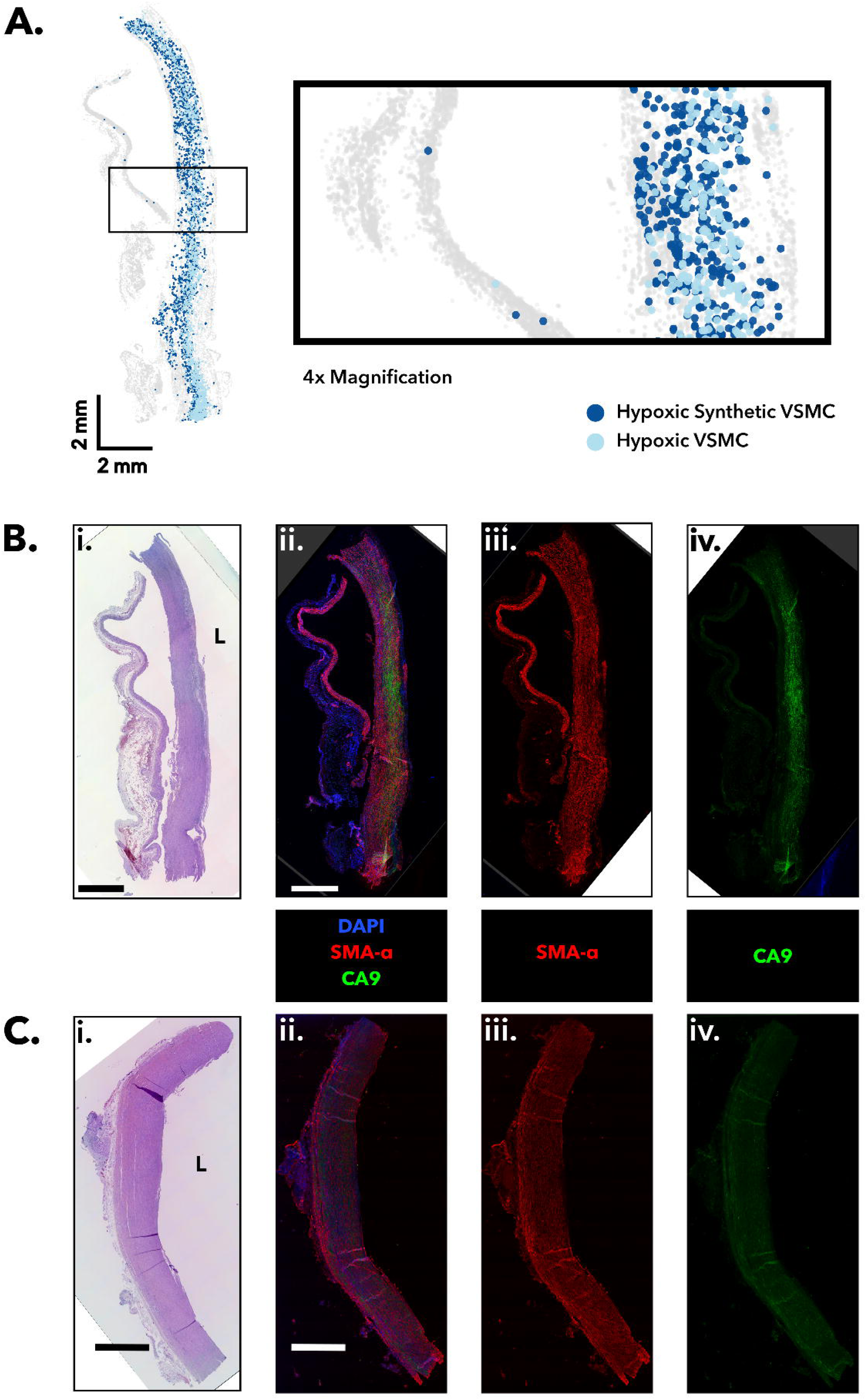
Pathologic and immunohistochemistry (IHC) validation of the presence of hypoxia-adapted vascular smooth muscle cells (VSMCs). **(A)** Spatial plot from a representative dissection tissue segment AAS 7.1 showing the distribution of hypoxic VSMC clusters. Box highlights 4x magnified region. Scale bars represent 2 mm. Plot displays adventitia to lumen from left to right. **(B)** Hematoxylin and Eosin (H&E) and immunohistochemistry (IHC) staining of the corresponding tissue dissection segment. **(i)** H&E. **(ii)** IHC labeling with 4′,6-diamidino-2-phenylindole (DAPI; blue), anti-smooth muscle actin-α (SMA-α; red), and anti-carbonic anhydrase 9 (CA9; green) demonstrating the presence of hypoxia markers in the mid-media. **(iii)** Single fluorescent channel (TRITC) image showing dropout of SMA-α in the corresponding region of increased CA9 intensity. **(iv)** Single fluorescent channel (FITC) image showing distribution of CA9 within the mid-media with highest intensity in the area of reduced SMA-α expression. (**C)** H&E and IHC of neighboring sections from a representative DC specimen (DC1). **(i)** H&E. **(ii)** IHC labeling with DAPI (blue), SMA-α (red), and CA9 (green). **(iii)** SMA-α only (red). **(iv)** CA9 only (green). Images highlight the relative absence of a hypoxic mid-medial core. L= lumen, DAPI = 4′,6-diamidino-2-phenylindole, SMA-α = smooth muscle actin-α, CA9 = carbonic anhydrase 9.

### VSMC spatial domains reveal distinct medial patterns in AAS and DC

To distinguish VSMC regions bordering the interlamellar separation or dissection tear from medial regions within the dissection flap, we applied BANKSY (λ=0.8) to AAS and DC aortic tissues. We identified six VSMC-enriched spatial domains in the AAS samples and three in DC specimens. This analysis revealed six unique VSMC domains among the AAS and three among DC samples (Fig. 4). The domain UMAPs show VSMC-enriched domains in color and all other domains in gray (Fig. 4A-B). Spatial maps of representative samples reveal a layered arrangement of domains across the aortic wall. In AAS samples (Fig. 4C-D), domains 2 and 7 define the boundaries of interlamellar separation. The remaining VSMC-enriched domains form striated bands across the medial lamellar units within the dissection flap, which account for most of the media. Domain 1 (dodger blue) contains hypoxia-adapted VSMC populations and does not directly border the tear. Spatial maps of representative DC specimens demonstrat that domain 2 predominates (Fig. 4D), although its transmural distribution varies among specimens. For example, it lies near the adventitial border in DC2 but closer to the luminal border in the other DC aortas.

**Figure 4:**
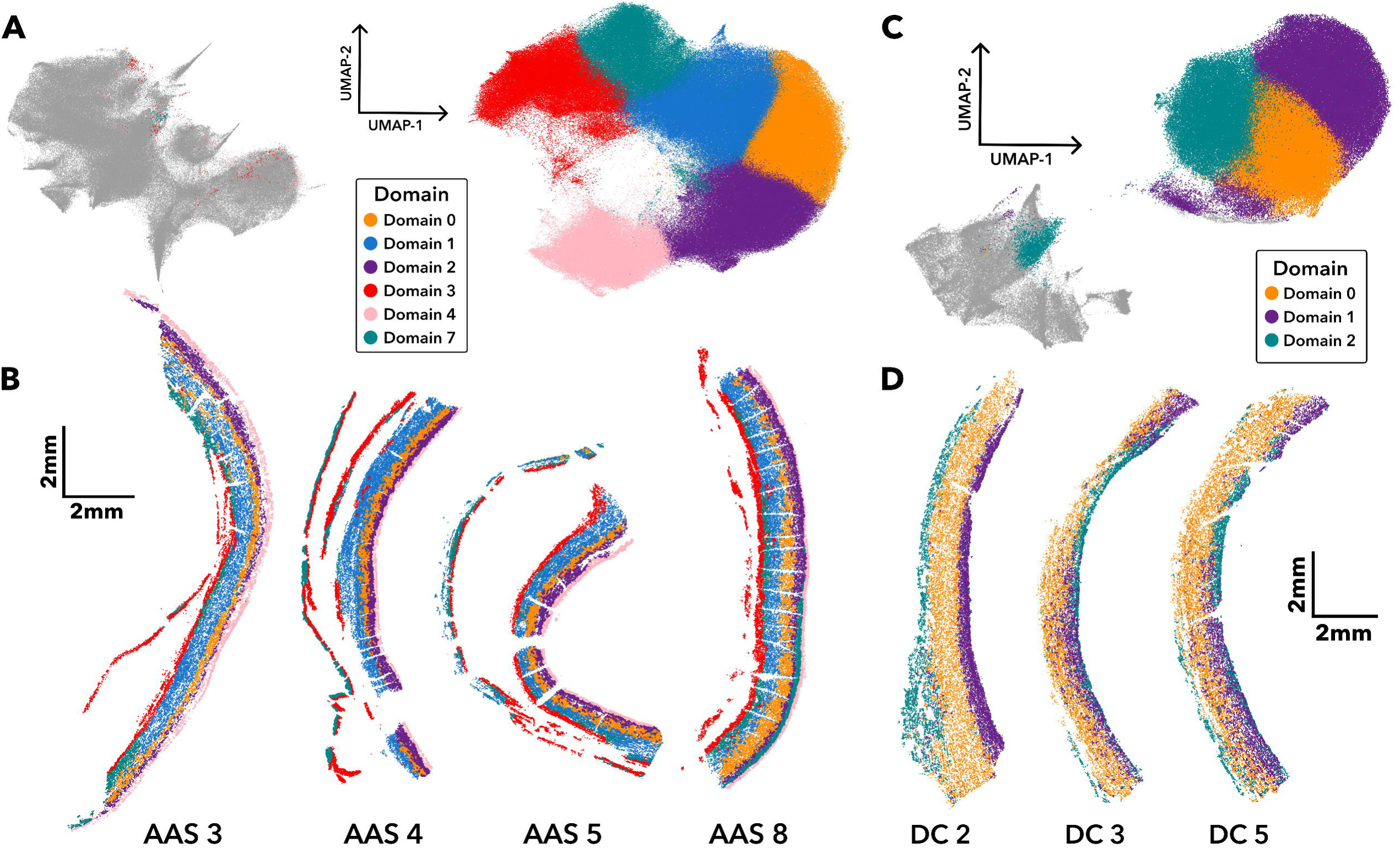
Domain mapping of Acute Aortic Syndrome (AAS) and Donor Control (DC) samples. (A) Uniform Manifold Projection (UMAP) of spatial domains of AAS samples. The right portion of the UMAP demonstrates the six VSMC spatial domains, whereas the left is the combined non-VSMC spatial domains. (B) Spatial plots of four representative AAS specimens showing the layered spatial orientation of each medial domain. Spatial plots display adventitia to lumen from left to right. (C) UMAP of spatial domains of DC samples. Similarly to AAS samples, the right portion of the UMAP contains the bulk of the cells within the three VSMC spatial domains, and the left is the combined non-VSMC spatial domains. (D) Spatial plots of three representative DC specimens showing layered spatial orientation of each medial domain. Spatial plots display adventitia to lumen from left to right.

### Striated VSMC transmural distribution in AAS but not in donor aortas

To quantify the positions of VSMC clusters across the layered aortic wall, we applied Laplace-based analysis with Dirichlet boundary conditions to AAS and DC specimens. DC samples were analyzed from intima to adventitia. Because of artificial spatial distances created during tissue processing, AAS specimens were analyzed from intima to dissection plane which includes the majority of the biologically relevant tissue. In AAS specimens, contractile and synthetic VSMC clusters occupied distinct depths (Fig. 5B). Inner Medial Contractile VSMCs were concentrated in the inner half of the media, whereas Outer Medial Contractile VSMCs were concentrated in the outer half. Similarly, Synthetic VSMCs were concentrated in the inner media, while Collagen-producing Synthetic VSMCs were most abundant in the outer media. Consistent with previous findings, hypoxia-adapted VSMC niches were concentrated near the mid-media.

**Figure 5:**
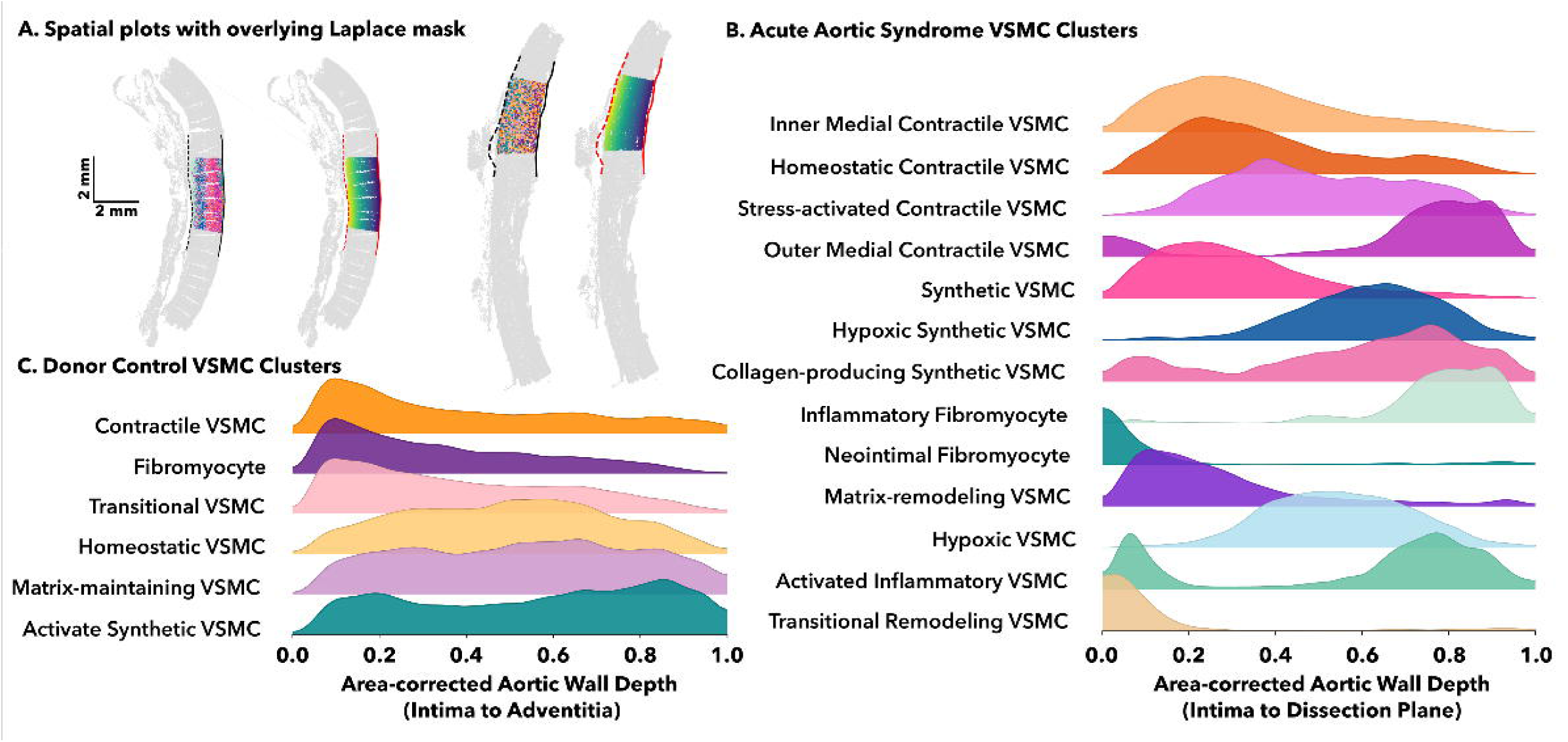
Laplace analysis of cellular depth distribution of VSMC clusters. **(A)** Representative spatial masks from Laplace analyses of AAS8 and DC1. Masks from full cohort found in Supplemental Figure 4A (AAS) and 5A (DC). Plots oriented from adventitia to lumen, left to right. **(B)** Area-corrected aortic wall depth of the 13 VSMC clusters present in the AAS samples. Depth is defined as intima=0 and dissection plane=1. **(C)** Area-corrected aortic wall depth of the 6 VSMC clusters present in the DC samples.

By contrast, DC aortic specimens had distinct depth distributions of VSMC niches (Fig. 5C). Contractile VSMCs, Fibromyocytes, and Transitional VSMCs were concentrated in the inner media. The remaining VSMC clusters had a more uniform depth distribution across the wall, a pattern not observed among VSMC clusters in AAS specimens.

## Discussion

### Spatial atlas and hypoxic remodeling

This study defines the first cellular-resolution spatial atlas of ascending aortic tissue in human AAS, revealing laminar patterns of VSMC phenotypic remodeling alongside hypoxia-associated adaptation. By mapping cell states within native tissue architecture, this approach resolves medial organization that is not readily captured by dissociation-based sc/sn RNA-sequencing (RNA-seq) methods and circumvents the challenges inherent in VSMC isolation.^22^ In AAS specimens, contractile, synthetic, and hypoxia-signaling VSMC clusters occupy distinct positions within the media, in contrast to the more uniform distributions of several VSMC clusters in DC aortas. Spatial domain analysis further distinguishes regions bordering the tear or interlamellar separation from the striated domains spanning the medial lamellae within the dissection flap. Among these, a recurrent hypoxia-adapted domain occupies the mid-media of the dissection flap, remains distinct from the tear-bordering regions, and corresponds to CA9-positive regions, with no comparable region in DC aortas. Together, these findings identify altered laminar organization and spatially localized hypoxia adaptation as complementary features of medial remodeling in AAS. They provide a cellular and spatial framework for investigating how these features may relate to medial vulnerability and for developing imaging strategies to assess their potential as markers of dissection risk.

Hypoxia and HIF-1α signaling have previously been implicated as important contributors to aortic dissection pathology. ^12,23^ In our spatial atlas, mid-medial hypoxia-signaling VSMCs showed marked transcriptional downregulation of contractile programs, a pattern mirrored at the protein level by reduced SMA-α immunoreactivity in the same regions. In vascular injury models, transcriptional downregulation of contractile genes requires sustained signaling over 24-72h, with medial VSMC contractile protein loss becoming apparent between days 3-7. These kinetics suggest that the observed remodeling developed over at least several days rather than within hours since dissection onset. Regions with reduced SMA-α protein expression also exhibited CA9 immunoreactivity, consistent with previous observations in human AAD. CA9 generally requires several hours of sustained, moderate hypoxia to become detectable, and its long protein half-life means makes it a marker of cumulative exposure rather than oxygen tension around the time of resection. Taken together with contractile downregulation, we interpret CA9 as evidence of sustained hypoxia-associated signaling. This distinction is relevant given the acute interval between symptom onset and surgical repair in our cohort (within 24 hours for 7/8 samples).^24^ Neither CA9 nor contractile-program loss alone establishes a chronic hypoxia-driven process, and CA9 can also be induced by non-hypoxic stimuli. Nevertheless, the spatial convergence of CA9 positivity with contractile-program loss at both the transcript and protein levels is unlikely to be explained solely by acute peri-dissection changes and instead supports the presence of a pre-existing hypoxia-associated, phenotypically modulated medial compartment. This organization argues against the interpretation that hypoxia-adapted VSMCs arise solely from vasa vasorum stripping during dissection. We suspect, instead, this region represents a chronic watershed hypoxic zone arising during hypertensive aortic remodeling, where medial lamellae are relatively distant from both luminal diffusion and vasa vasorum-derived oxygen and nutrient delivery.

### Xenium resolution compared with sc/snRNA-seq

Our use of Xenium in situ hybridization enabled profiling across large aortic segments using a custom panel designed primarily to resolve VSMC phenotypic states. This approach identified 14 transcriptionally defined VSMC states in their native spatial context in AAS specimens. Compared with dissociation-based sc/snRNA-seq, in situ analysis preserves native spatial architecture, enabling direct alignment with transmural tissue depth and relationships to regions of tissue cleavage. It additionally circumvents the significant under sampling of VSMCs relative to other rarer cell types shown in multiple previous investigations, presumptively related to challenges in VSMC isolation.^22,26^ Within this framework, the 14 states fall into familiar major categories—contractile, synthetic, inflammatory, and hypoxia-adapted—but are further refined by lamellar depth. Profiling large aortic segments spatially provides substantially greater cell counts compared with studies of thoracic aorta, increasing power to detect rare states and distinguish closely related phenotypes occupying distinct spatial niches. Together, these features highlight the complementary strengths of Xenium in situ hybridization relative to dissociation-based methods and support the biological basis for the spatially organized VSMC heterogeneity we observe in AAS.

### Limitations and future work

A key limitation of this study is that human AAS specimens cannot establish causality nor define the temporal sequence by which medial remodeling develops before dissection. Also, we cannot definitively determine what transcriptional states are altered in the setting of acute interlamellar separation with AAS and in the hours between dissection onset and repair. We therefore cannot determine whether transmural medial striation or the mid-medial hypoxia-associated VSMC domain definitively predispose to interlamellar separation and dissection.

Nevertheless, examining human AAS tissue differs fundamentally from settings in which tissue is sampled only at the terminal stage of progressive organ failure. We interpret sporadic AAS as arising when an acute hemodynamic trigger acts on a variably remodeled, permissive substrate. Because this substrate does not itself determine when—or whether—dissection occurs, the aortic wall we sample reflects the predisposing background rather than a fixed end stage of unavoidable failure. The severity of this background likely varies between cases, such that a similar trigger may or may not precipitate tearing, depending on the extent of underlying remodeling. Although our findings are correlative, the dissected wall therefore likely captures features of the substrate that enables dissection, with some superimposed acute changes, rather than only the terminal consequences of a chronic disease process. The consistent identification of increased laminar remodeling and a hypoxia-adapted domain in AAS, but not donor aortas, supports its relevance to the medial remodeling landscape associated with dissection.

Additionally, this given that Xenium requires segmentation of the human aorta given the area available for imaging and the sampling design of this study, these findings are more reflective of the transmural cellular architecture of the lesser curvature in AAS and DC ascending aortic tissues. We thought examining the intact wall and the dissection margin would be particularly interesting for understanding the cleavage plane and its termination. Therefore, further studies will be needed to examine the human aortic wall circumferentially, given the differential aortic stress in the greater and lesser curvatures. This will help us better understand remodeling due to hemodynamic stress in the ascending aorta.

These data suggest several paths for future work. Future studies should determine how spatially distinct VSMC niches shape local extracellular matrix composition and organization, and whether regional downregulation of contractile programs produces corresponding differences in medial distensibility across the aortic wall. Integrating spatial transcriptomics with proteomic, biomechanical, and high-resolution matrix imaging approaches may help define whether these molecular and structural gradients create focal susceptibility to interlamellar separation.

Additional studies will be required to define the temporal sequence of disease progression changes. Animal models may be informative, but if medial hypoxia is a precondition for dissection, relevant systems will need to recapitulate the human hypoxic medial environment, which may limit the utility of thin-walled rodent models. This study specifically examines aortas from AAS patients with no syndromic features or suspected history of familial connective tissue disorders, though we cannot rule out undiagnosed connective tissue abnormalities in our cohort. However, future studies applying spatial transcriptomic approaches to cohorts with genetically defined connective tissue disorders will be important for determining whether the spatial patterns of VSMC phenotypic modulation observed in our patient with Marfan syndrome are reproducible and shared across heritable aortopathies.

AI disclosure: During preparation of this work the author(s) used Perplexity.ai in order to assist with cluster identification and editing of text for clarity. After using this tool/service, the author(s) reviewed and edited the content as needed and take(s) full responsibility for the content of the publication.

## Abbreviations

AAS: Acute aortic syndrome
ATAAD: Acute type A aortic dissection
ATAIMH: Acute type A intramural hematoma
BANKSY: Building Aggregates with a Neighborhood Kernel and Spatial Yardstick
BSA: Bovine Serum Albumin
C: Celsius
DAPI: 4′,6-diamidino-2-phenylindole
DC: Donor control
HTN: Hypertension
H&E: Hematoxylin and eosin
IHC: Immunohistochemistry
RNA: Ribonucleic acid
sc: Single-cell
sn: Single-nucleus
TBST: Tris-Buffered Saline with Tween 20
UMAP: Uniform Manifold Projection
VSMC: Vascular smooth muscle cell

