## Supplemental Methods pt 1 for "Cellular-Resolution Spatial Transcriptomics Reveals Laminar VSMC Phenotypic Remodeling and a Hypoxic Medial Core in Human Thoracic Aortic Dissection"

#### BANKSY λ comparison — Leiden(res=1.2) UMAP (aligned to λ=0.2)

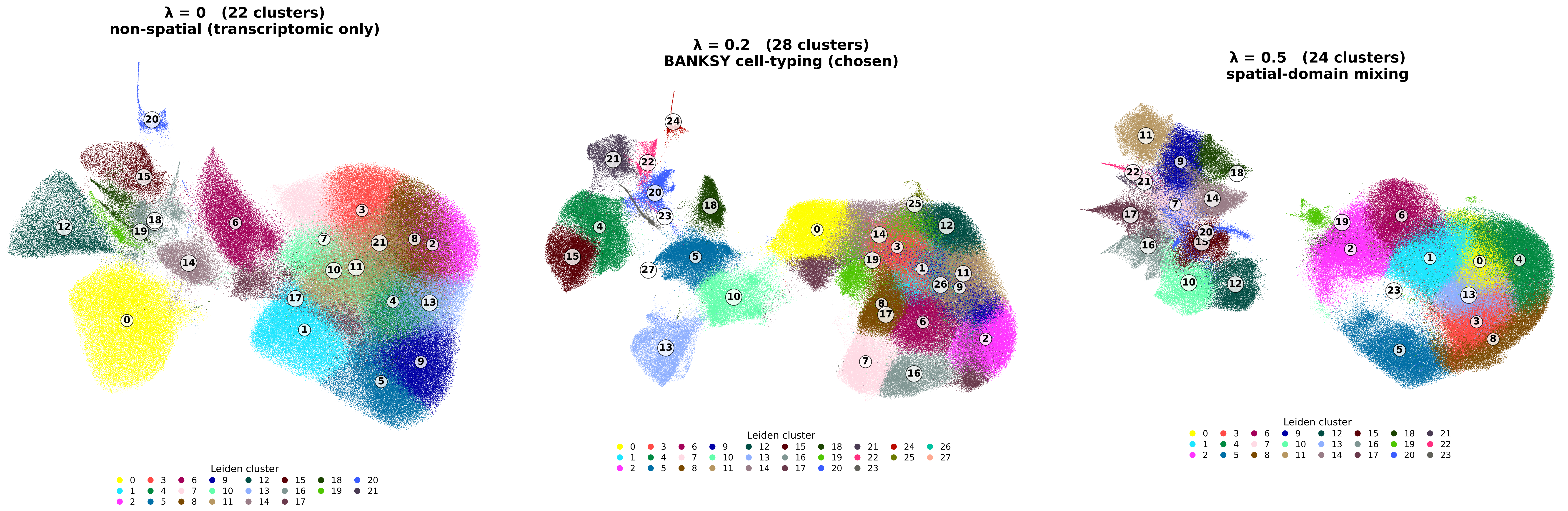

**Figure 10c — BANKSY PCA (Unintegrated) vs Harmony Integration: UMAP & Silhouette**
**(UMAP: n = 80,000 cells subsampled, colored by sample)**

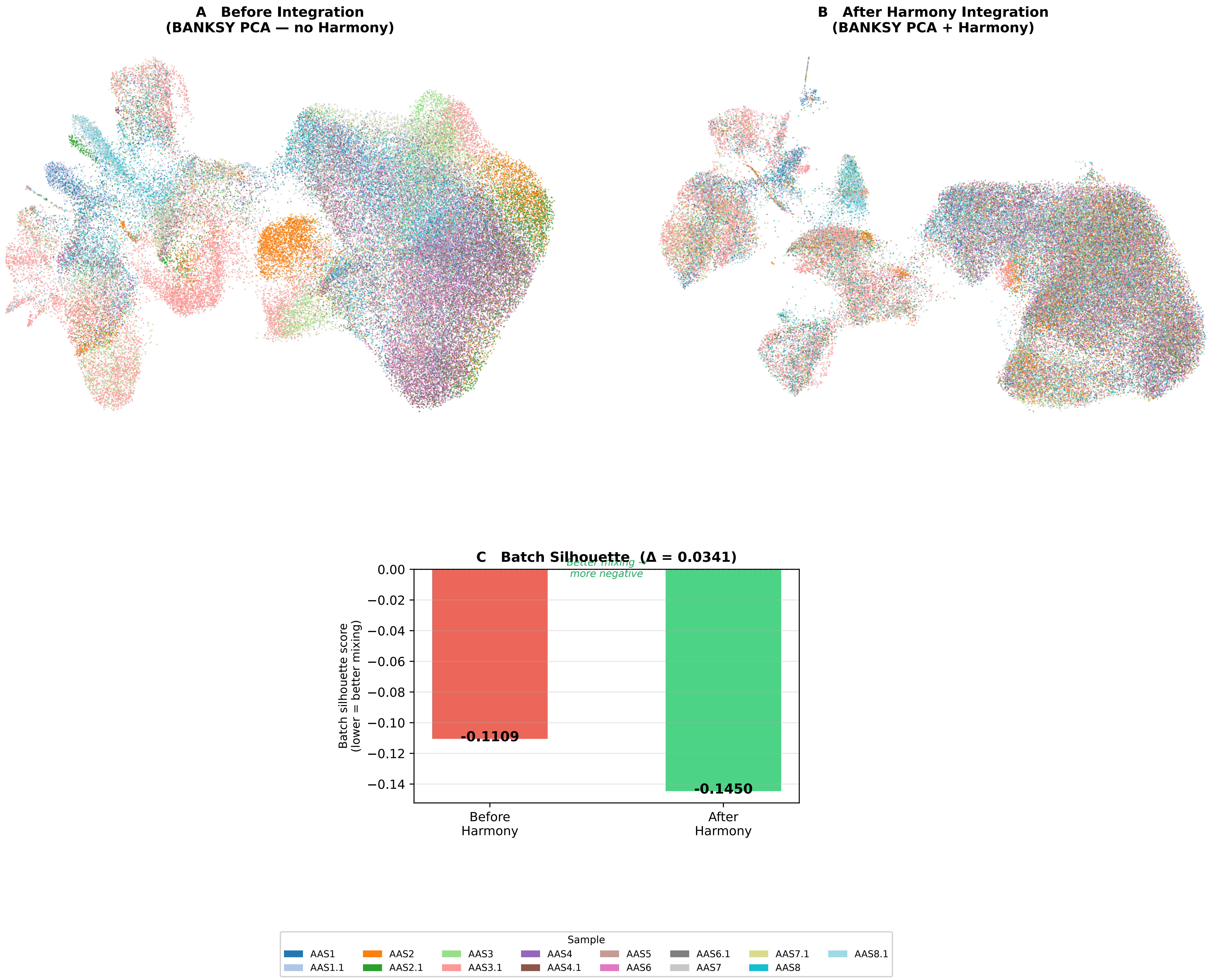

**Figure 8 — Harmony Batch Correction: Pre vs. Post UMAP**
**(n = 80,000 cells, colored by sample)**

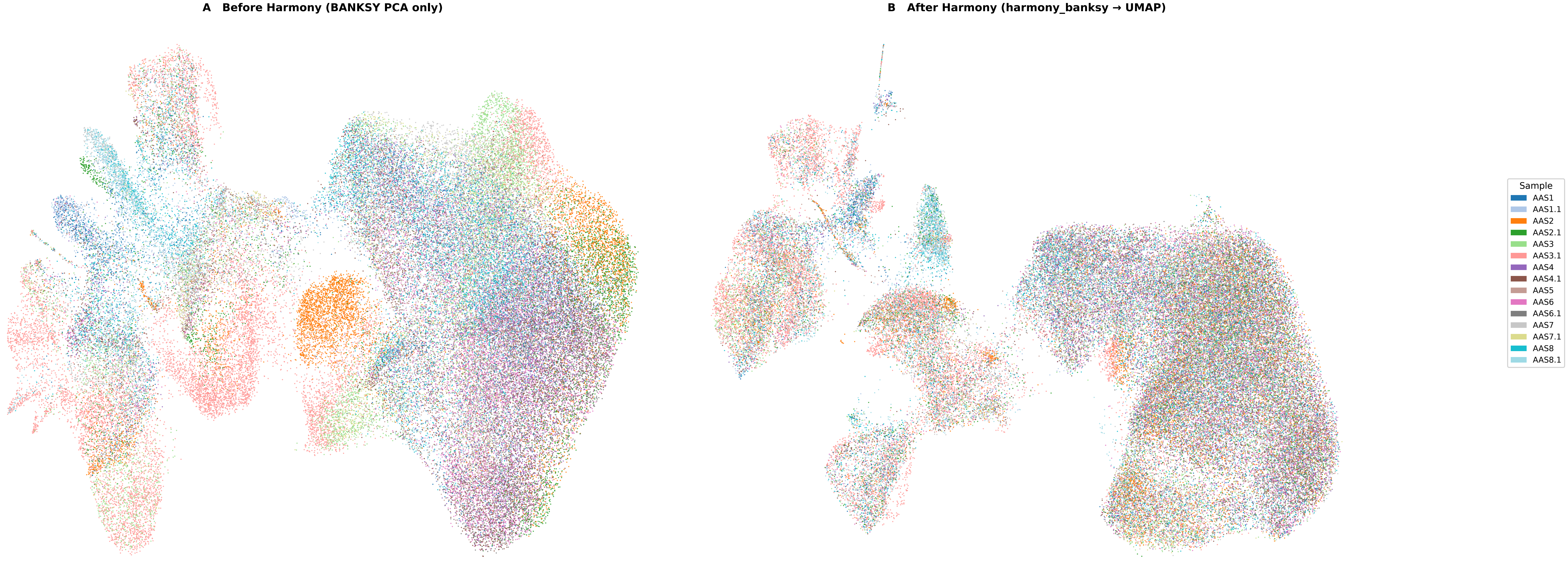

**Figure 10b — LISI Validation: Harmony Batch Mixing & Cluster Purity**
**iLISI median = 8.82 (PASS) | cLISI median = 1.27 (PASS) | n = 99,990 cells (stratified subsample)**

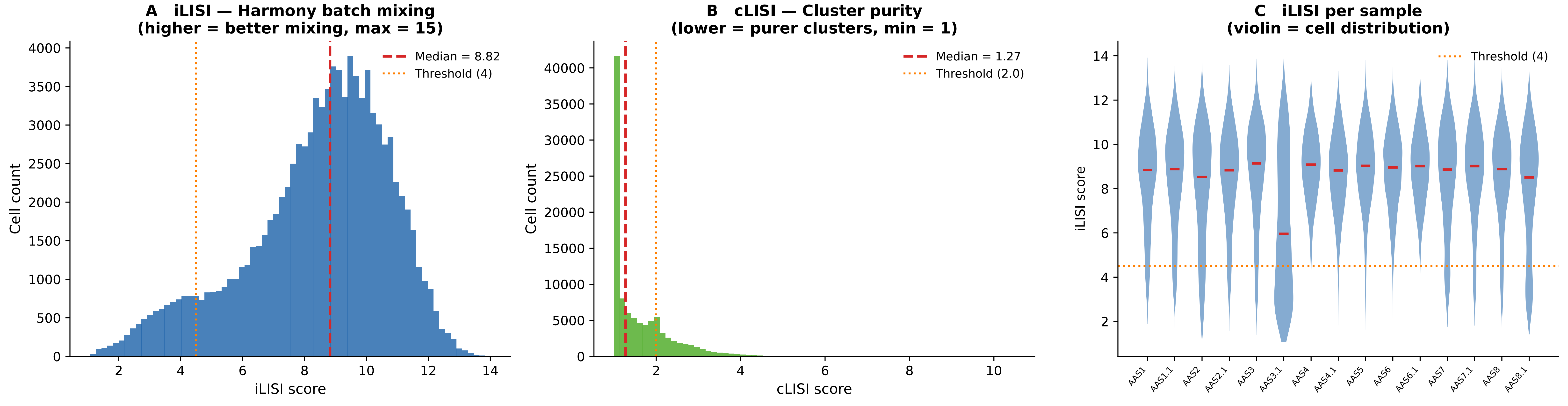

### Leiden resolution sweep (λ=0.2, n50 euclidean UMAP)

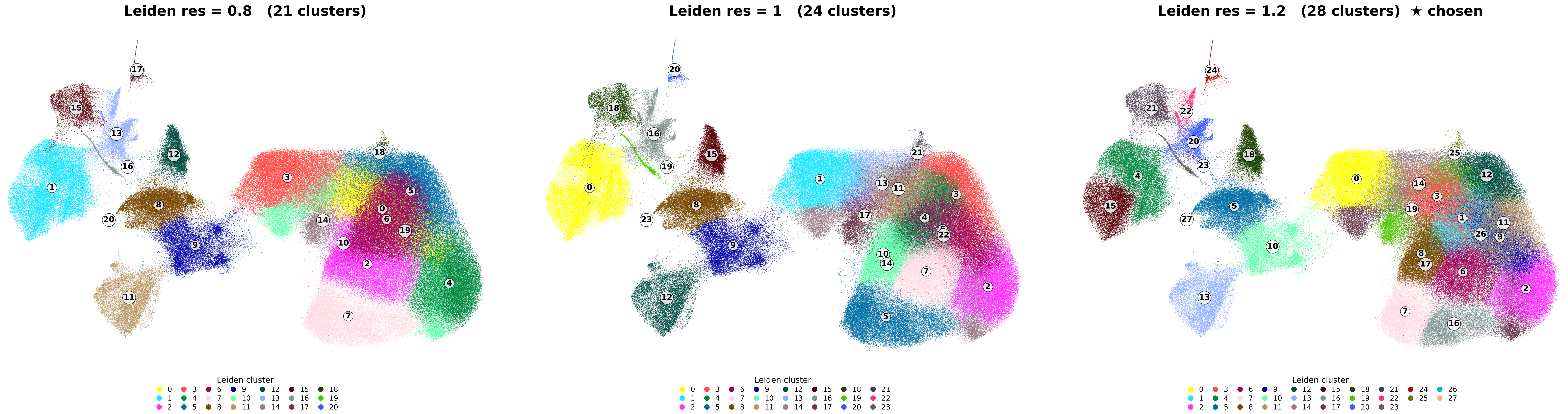
