## Supplemental Methods pt 2 for "Cellular-Resolution Spatial Transcriptomics Reveals Laminar VSMC Phenotypic Remodeling and a Hypoxic Medial Core in Human Thoracic Aortic Dissection"

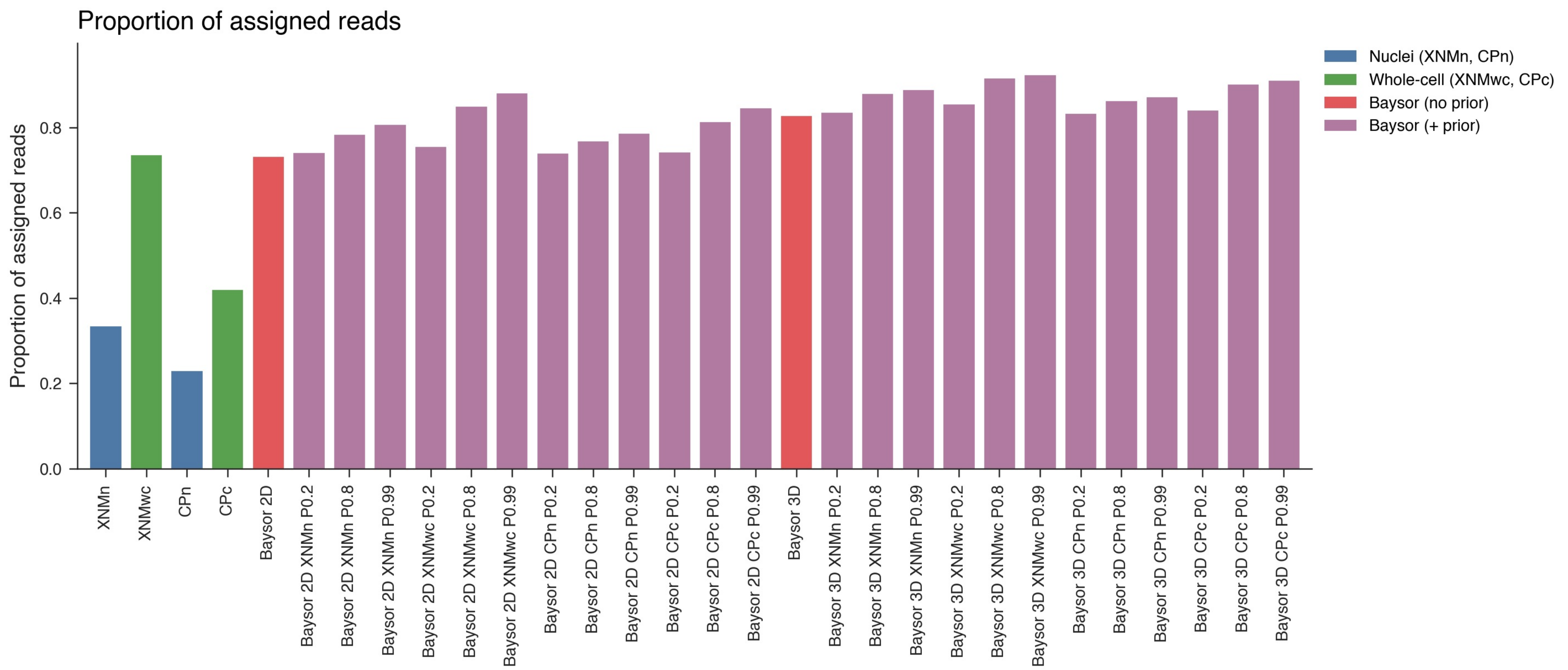

Proportion of assigned reads
Proportion of assigned reads
0.0
0.2
0.4
0.6
0.8
Nuclei (XNMn, CPn)
Whole-cell (XNMwc, CPc)
Baysor (no prior)
Baysor (+ prior)
XNMn
XNMwc
CPn
CPc
Baysor 2D
Baysor 2D XNMn P0.2
Baysor 2D XNMn P0.8
Baysor 2D XNMn P0.99
Baysor 2D XNMwc P0.2
Baysor 2D XNMwc P0.8
Baysor 2D XNMwc P0.99
Baysor 2D CPn P0.2
Baysor 2D CPn P0.8
Baysor 2D CPn P0.99
Baysor 2D CPc P0.2
Baysor 2D CPc P0.8
Baysor 2D CPc P0.99
Baysor 3D
Baysor 3D XNMn P0.2
Baysor 3D XNMn P0.8
Baysor 3D XNMn P0.99
Baysor 3D XNMwc P0.2
Baysor 3D XNMwc P0.8
Baysor 3D XNMwc P0.99
Baysor 3D CPn P0.2
Baysor 3D CPn P0.8
Baysor 3D CPn P0.99
Baysor 3D CPc P0.2
Baysor 3D CPc P0.8
Baysor 3D CPc P0.99

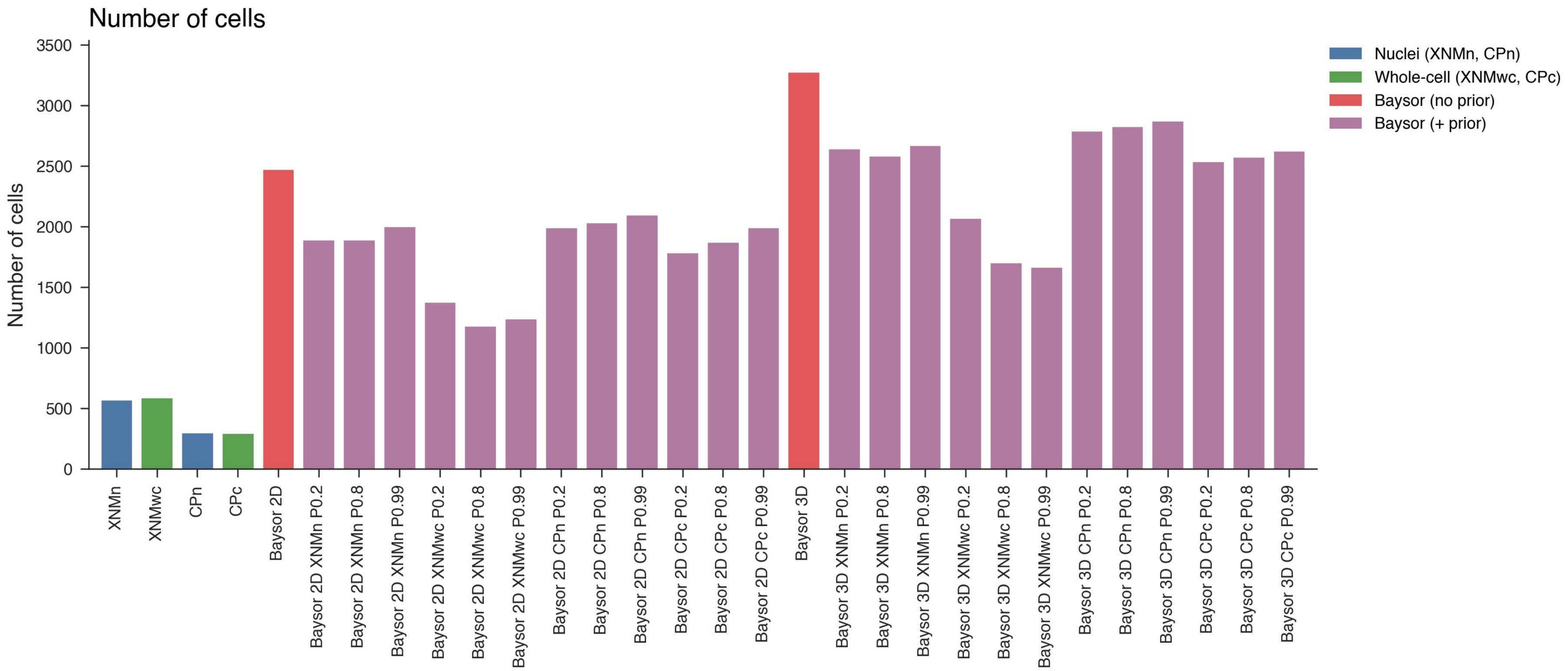

Number of cells
Number of cells
0
500
1000
1500
2000
2500
3000
3500
Nuclei (XNMn, CPn)
Whole-cell (XNMwc, CPc)
Baysor (no prior)
Baysor (+ prior)
XNMn
XNMwc
CPn
CPc
Baysor 2D
Baysor 2D XNMn P0.2
Baysor 2D XNMn P0.8
Baysor 2D XNMn P0.99
Baysor 2D XNMwc P0.2
Baysor 2D XNMwc P0.8
Baysor 2D XNMwc P0.99
Baysor 2D CPn P0.2
Baysor 2D CPn P0.8
Baysor 2D CPn P0.99
Baysor 2D CPc P0.2
Baysor 2D CPc P0.8
Baysor 2D CPc P0.99
Baysor 3D
Baysor 3D XNMn P0.2
Baysor 3D XNMn P0.8
Baysor 3D XNMn P0.99
Baysor 3D XNMwc P0.2
Baysor 3D XNMwc P0.8
Baysor 3D XNMwc P0.99
Baysor 3D CPn P0.2
Baysor 3D CPn P0.8
Baysor 3D CPn P0.99
Baysor 3D CPc P0.2
Baysor 3D CPc P0.8
Baysor 3D CPc P0.99

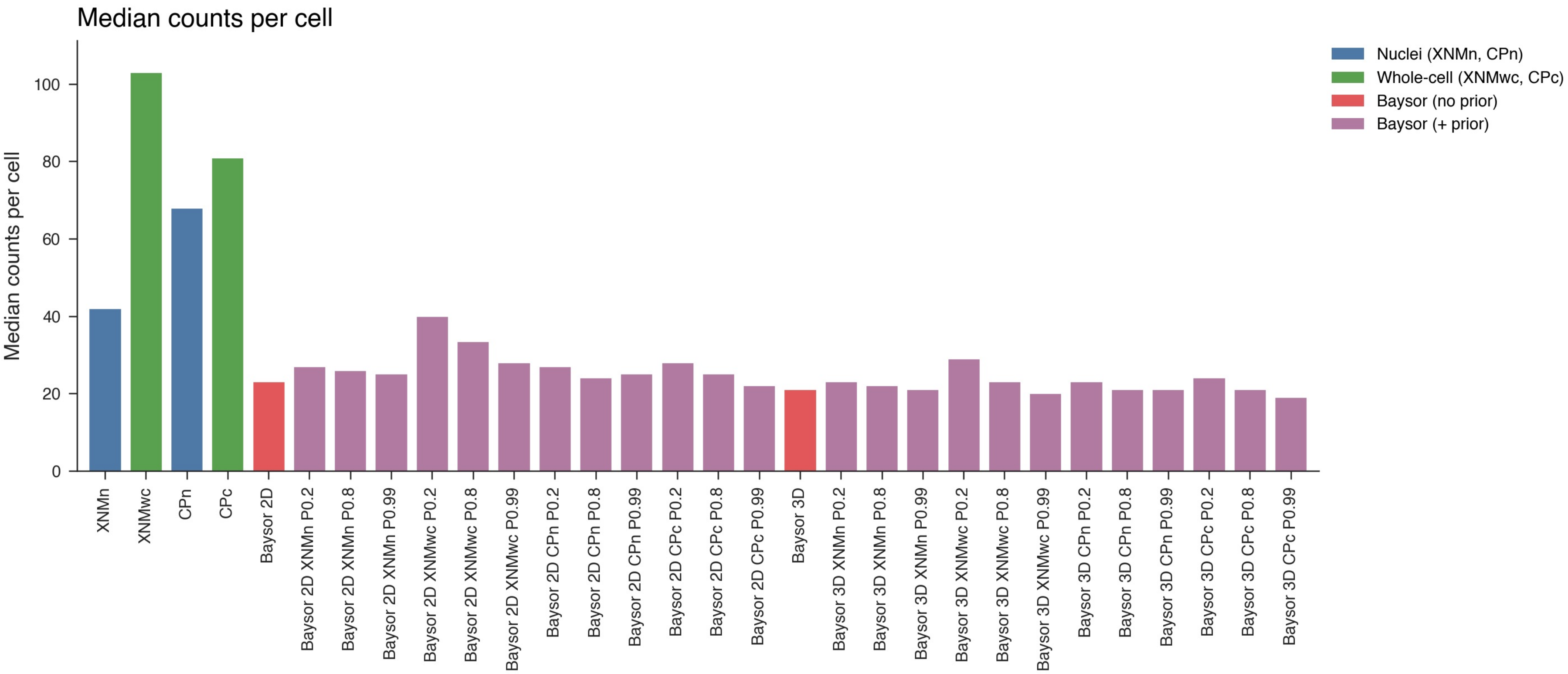
Median counts per cell
Median counts per cell
0
20
40
60
80
100
Nuclei (XNMn, CPn)
Whole-cell (XNMwc, CPc)
Baysor (no prior)
Baysor (+ prior)
XNMn
XNMwc
CPn
CPc
Baysor 2D
Baysor 2D XNMn P0.2
Baysor 2D XNMn P0.8
Baysor 2D XNMn P0.99
Baysor 2D XNMwc P0.2
Baysor 2D XNMwc P0.8
Baysor 2D XNMwc P0.99
Baysor 2D CPn P0.2
Baysor 2D CPn P0.8
Baysor 2D CPn P0.99
Baysor 2D CPc P0.2
Baysor 2D CPc P0.8
Baysor 2D CPc P0.99
Baysor 3D
Baysor 3D XNMn P0.2
Baysor 3D XNMn P0.8
Baysor 3D XNMn P0.99
Baysor 3D XNMwc P0.2
Baysor 3D XNMwc P0.8
Baysor 3D XNMwc P0.99
Baysor 3D CPn P0.2
Baysor 3D CPn P0.8
Baysor 3D CPn P0.99
Baysor 3D CPc P0.2
Baysor 3D CPc P0.8
Baysor 3D CPc P0.99

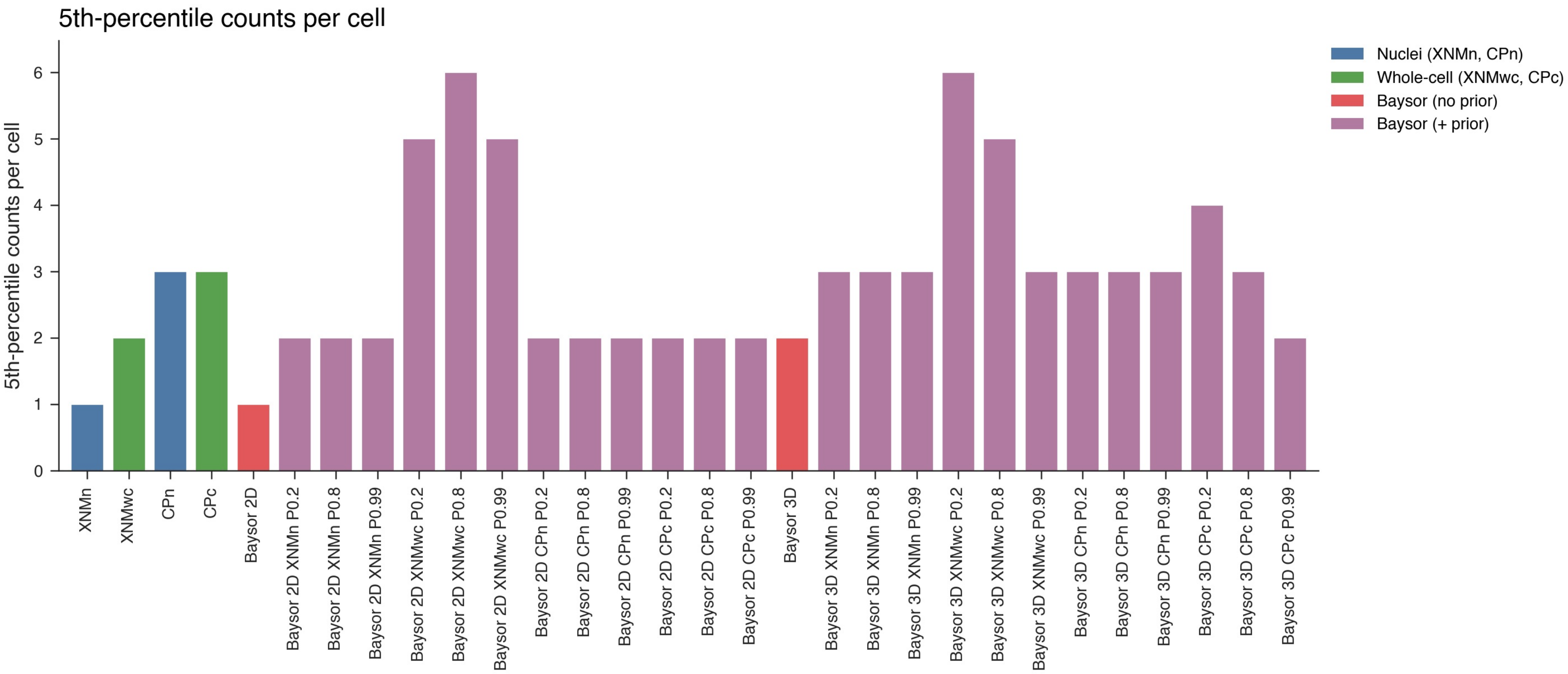
5th-percentile counts per cell
5th-percentile counts per cell
0
1
2
3
4
5
6
Nuclei (XNMn, CPn)
Whole-cell (XNMwc, CPc)
Baysor (no prior)
Baysor (+ prior)
XNMn
XNMwc
CPn
CPc
Baysor 2D
Baysor 2D XNMn P0.2
Baysor 2D XNMn P0.8
Baysor 2D XNMn P0.99
Baysor 2D XNMwc P0.2
Baysor 2D XNMwc P0.8
Baysor 2D XNMwc P0.99
Baysor 2D CPn P0.2
Baysor 2D CPn P0.8
Baysor 2D CPn P0.99
Baysor 2D CPc P0.2
Baysor 2D CPc P0.8
Baysor 2D CPc P0.99
Baysor 3D
Baysor 3D XNMn P0.2
Baysor 3D XNMn P0.8
Baysor 3D XNMn P0.99
Baysor 3D XNMwc P0.2
Baysor 3D XNMwc P0.8
Baysor 3D XNMwc P0.99
Baysor 3D CPn P0.2
Baysor 3D CPn P0.8
Baysor 3D CPn P0.99
Baysor 3D CPc P0.2
Baysor 3D CPc P0.8
Baysor 3D CPc P0.99

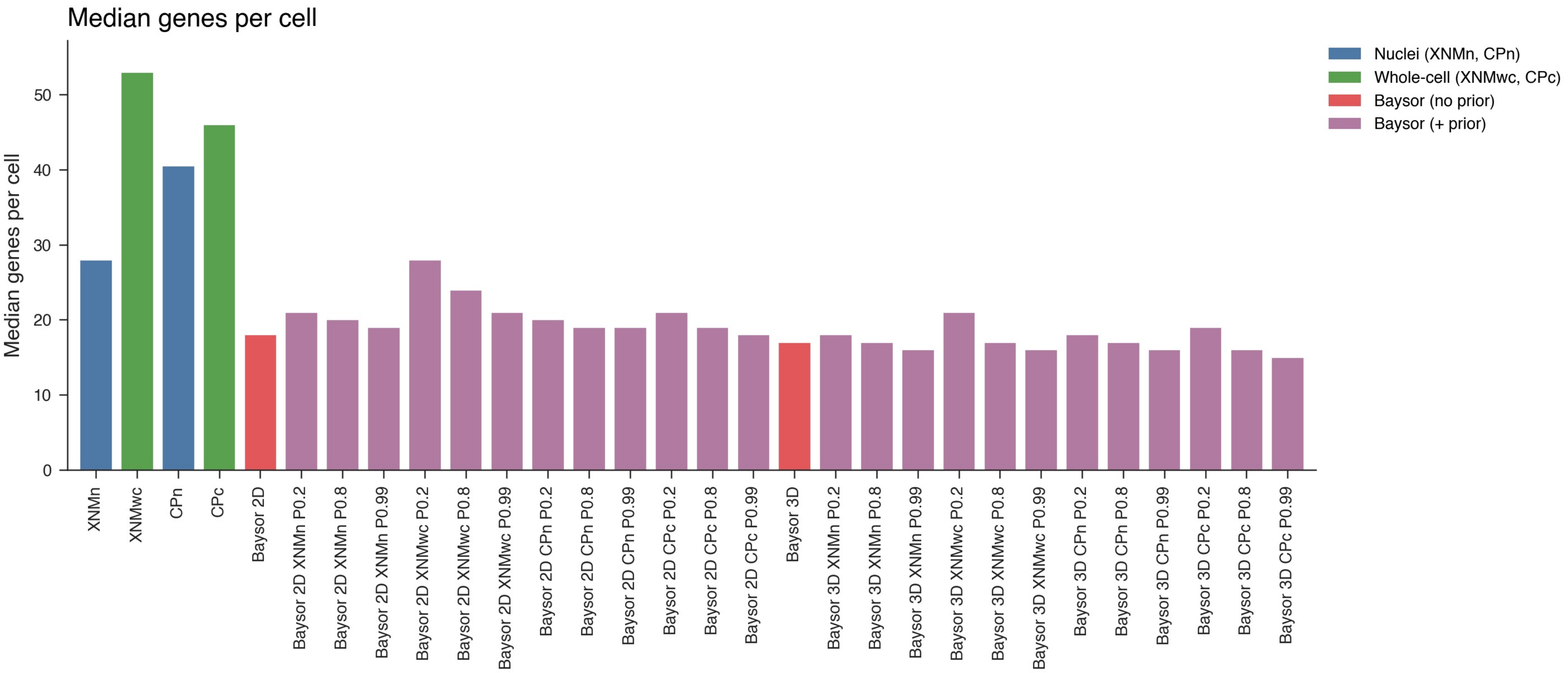
Median genes per cell
Median genes per cell
0
10
20
30
40
50
Nuclei (XNMn, CPn)
Whole-cell (XNMwc, CPc)
Baysor (no prior)
Baysor (+ prior)
XNMn
XNMwc
CPn
CPc
Baysor 2D
Baysor 2D XNMn P0.2
Baysor 2D XNMn P0.8
Baysor 2D XNMn P0.99
Baysor 2D XNMwc P0.2
Baysor 2D XNMwc P0.8
Baysor 2D XNMwc P0.99
Baysor 2D CPn P0.2
Baysor 2D CPn P0.8
Baysor 2D CPn P0.99
Baysor 2D CPc P0.2
Baysor 2D CPc P0.8
Baysor 2D CPc P0.99
Baysor 3D
Baysor 3D XNMn P0.2
Baysor 3D XNMn P0.8
Baysor 3D XNMn P0.99
Baysor 3D XNMwc P0.2
Baysor 3D XNMwc P0.8
Baysor 3D XNMwc P0.99
Baysor 3D CPn P0.2
Baysor 3D CPn P0.8
Baysor 3D CPn P0.99
Baysor 3D CPc P0.2
Baysor 3D CPc P0.8
Baysor 3D CPc P0.99

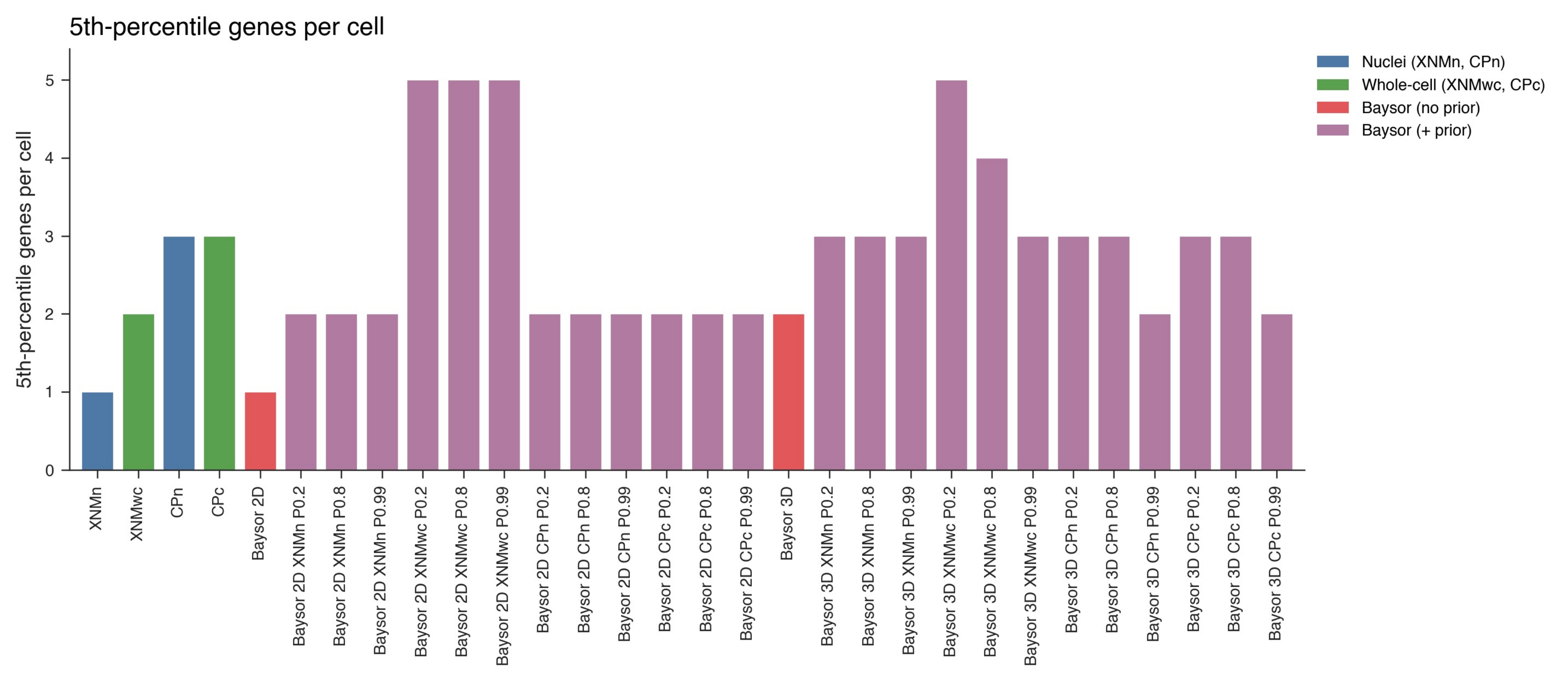
5th-percentile genes per cell
5th-percentile genes per cell
Nuclei (XNMn, CPn)
Whole-cell (XNMwc, CPc)
Baysor (no prior)
Baysor (+ prior)
0
1
2
3
4
5
XNMn
XNMwc
CPn
CPc
Baysor 2D
Baysor 2D XNMn P0.2
Baysor 2D XNMn P0.8
Baysor 2D XNMn P0.99
Baysor 2D XNMwc P0.2
Baysor 2D XNMwc P0.8
Baysor 2D XNMwc P0.99
Baysor 2D CPn P0.2
Baysor 2D CPn P0.8
Baysor 2D CPn P0.99
Baysor 2D CPc P0.2
Baysor 2D CPc P0.8
Baysor 2D CPc P0.99
Baysor 3D
Baysor 3D XNMn P0.2
Baysor 3D XNMn P0.8
Baysor 3D XNMn P0.99
Baysor 3D XNMwc P0.2
Baysor 3D XNMwc P0.8
Baysor 3D XNMwc P0.99
Baysor 3D CPn P0.2
Baysor 3D CPn P0.8
Baysor 3D CPn P0.99
Baysor 3D CPc P0.2
Baysor 3D CPc P0.8
Baysor 3D CPc P0.99

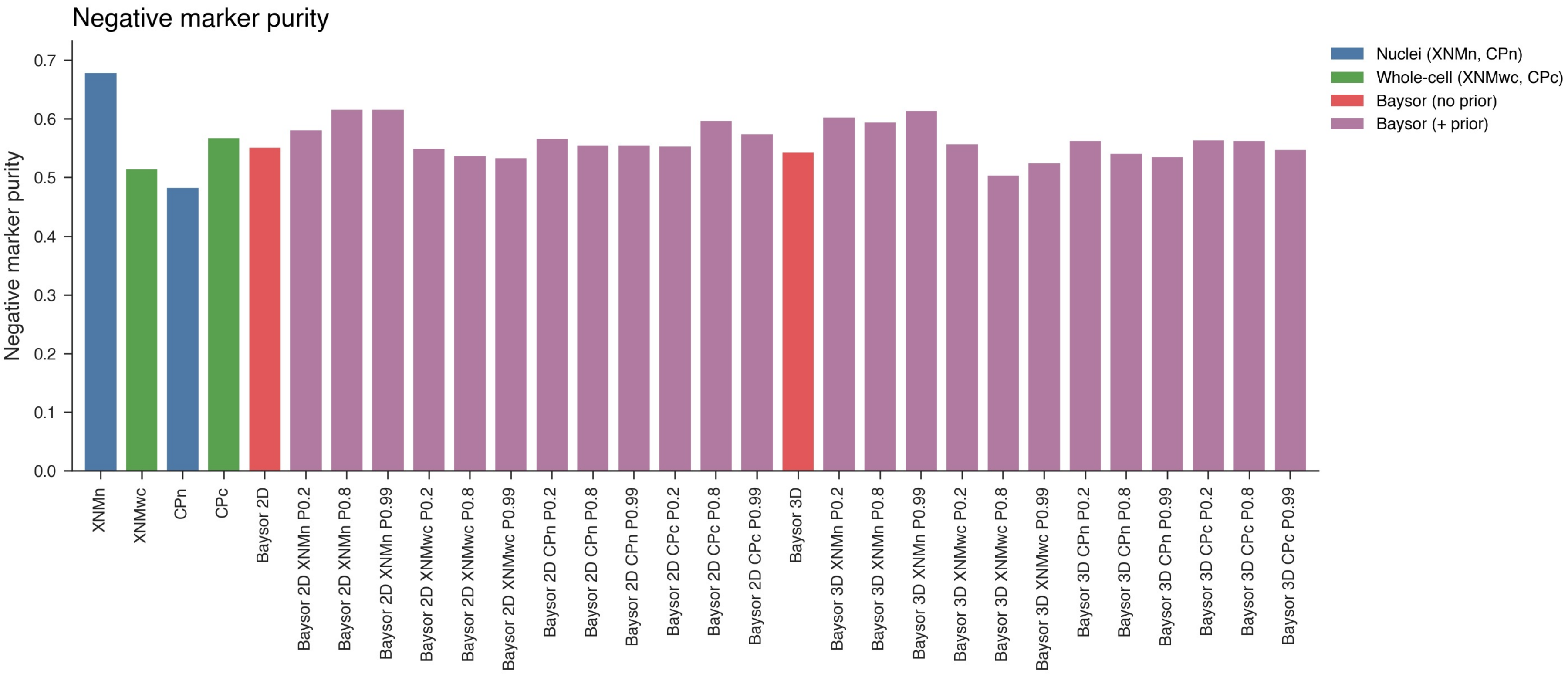
Negative marker purity
Negative marker purity
0.0
0.1
0.2
0.3
0.4
0.5
0.6
0.7
Nuclei (XNMn, CPn)
Whole-cell (XNMwc, CPc)
Baysor (no prior)
Baysor (+ prior)
XNMn
XNMwc
CPn
CPc
Baysor 2D
Baysor 2D XNMn P0.2
Baysor 2D XNMn P0.8
Baysor 2D XNMn P0.99
Baysor 2D XNMwc P0.2
Baysor 2D XNMwc P0.8
Baysor 2D XNMwc P0.99
Baysor 2D CPn P0.2
Baysor 2D CPn P0.8
Baysor 2D CPn P0.99
Baysor 2D CPc P0.2
Baysor 2D CPc P0.8
Baysor 2D CPc P0.99
Baysor 3D
Baysor 3D XNMn P0.2
Baysor 3D XNMn P0.8
Baysor 3D XNMn P0.99
Baysor 3D XNMwc P0.2
Baysor 3D XNMwc P0.8
Baysor 3D XNMwc P0.99
Baysor 3D CPn P0.2
Baysor 3D CPn P0.8
Baysor 3D CPn P0.99
Baysor 3D CPc P0.2
Baysor 3D CPc P0.8
Baysor 3D CPc P0.99

Segmentation metric summary

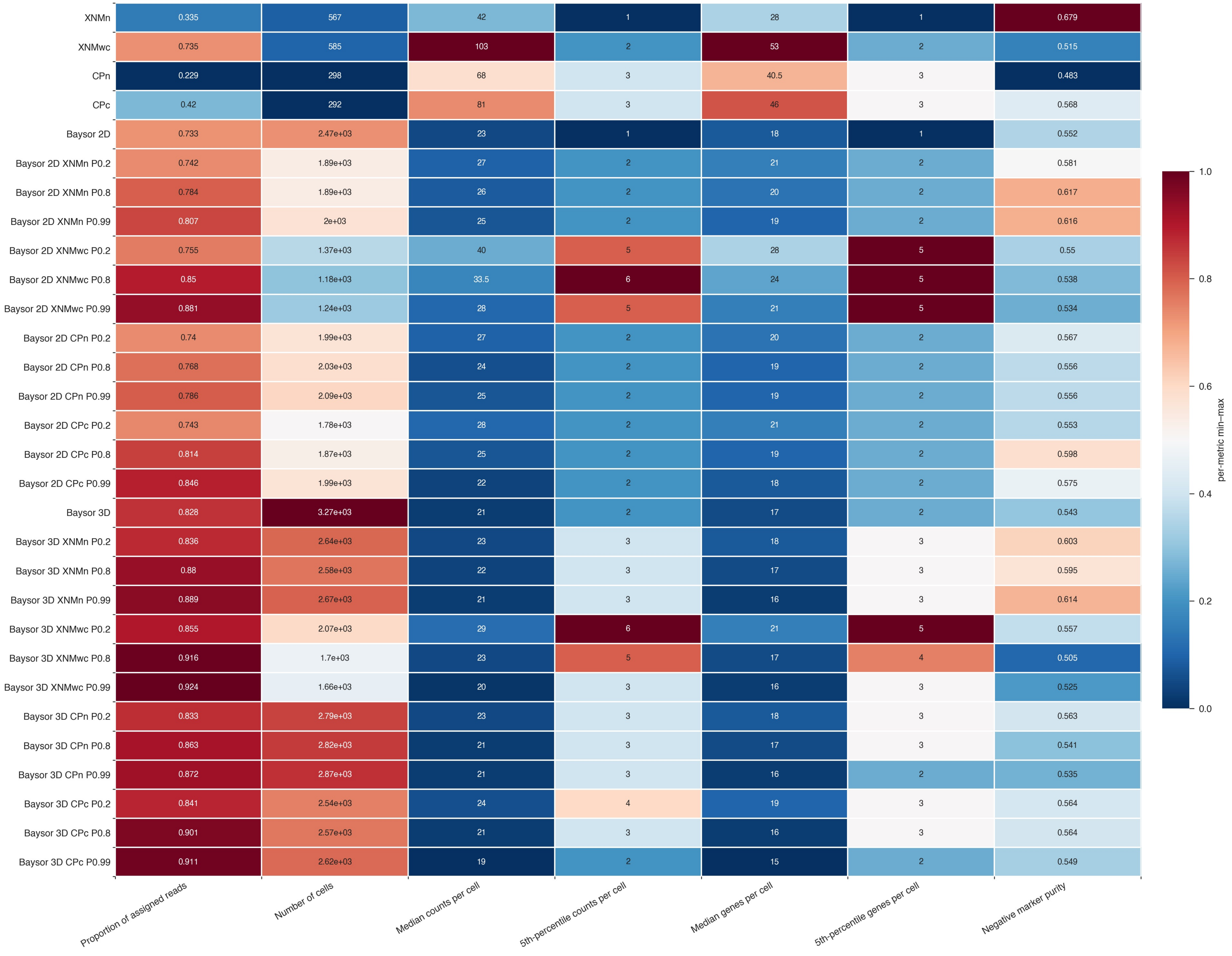

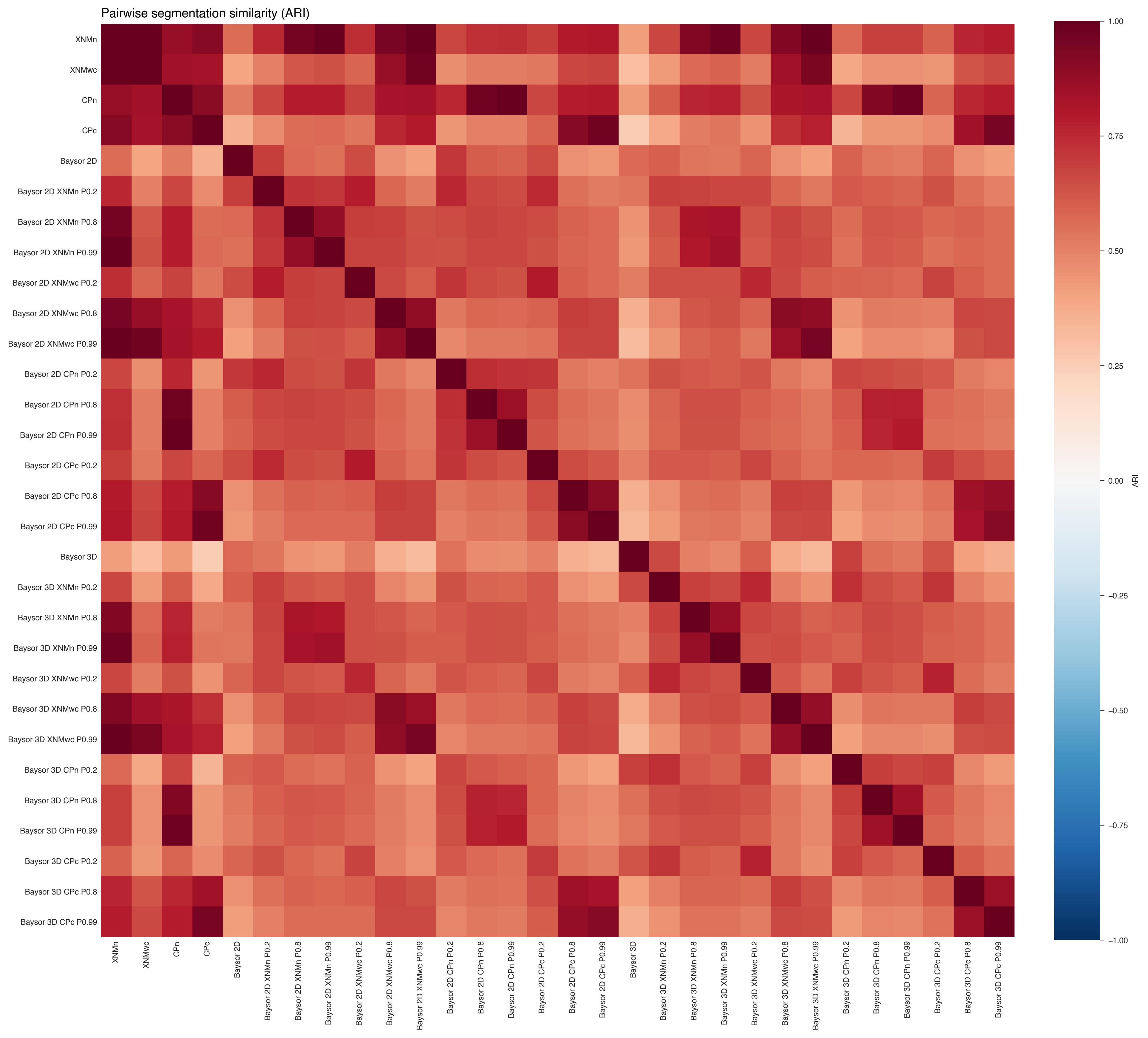
Pairwise segmentation similarity (ARI)
XNMn
XNMwc
CPn
CPc
Baysor 2D
Baysor 2D XNMn P0.2
Baysor 2D XNMn P0.8
Baysor 2D XNMn P0.99
Baysor 2D XNMwc P0.2
Baysor 2D XNMwc P0.8
Baysor 2D XNMwc P0.99
Baysor 2D CPn P0.2
Baysor 2D CPn P0.8
Baysor 2D CPn P0.99
Baysor 2D CPc P0.2
Baysor 2D CPc P0.8
Baysor 2D CPc P0.99
Baysor 3D
Baysor 3D XNMn P0.2
Baysor 3D XNMn P0.8
Baysor 3D XNMn P0.99
Baysor 3D XNMwc P0.2
Baysor 3D XNMwc P0.8
Baysor 3D XNMwc P0.99
Baysor 3D CPn P0.2
Baysor 3D CPn P0.8
Baysor 3D CPn P0.99
Baysor 3D CPc P0.2
Baysor 3D CPc P0.8
Baysor 3D CPc P0.99
ARI
1.00
0.75
0.50
0.25
0.00
−0.25
−0.50
−0.75
−1.00

**Figure 4c — Segmentation QC: Full-Tissue Overview and DAPI Zoom Validation**

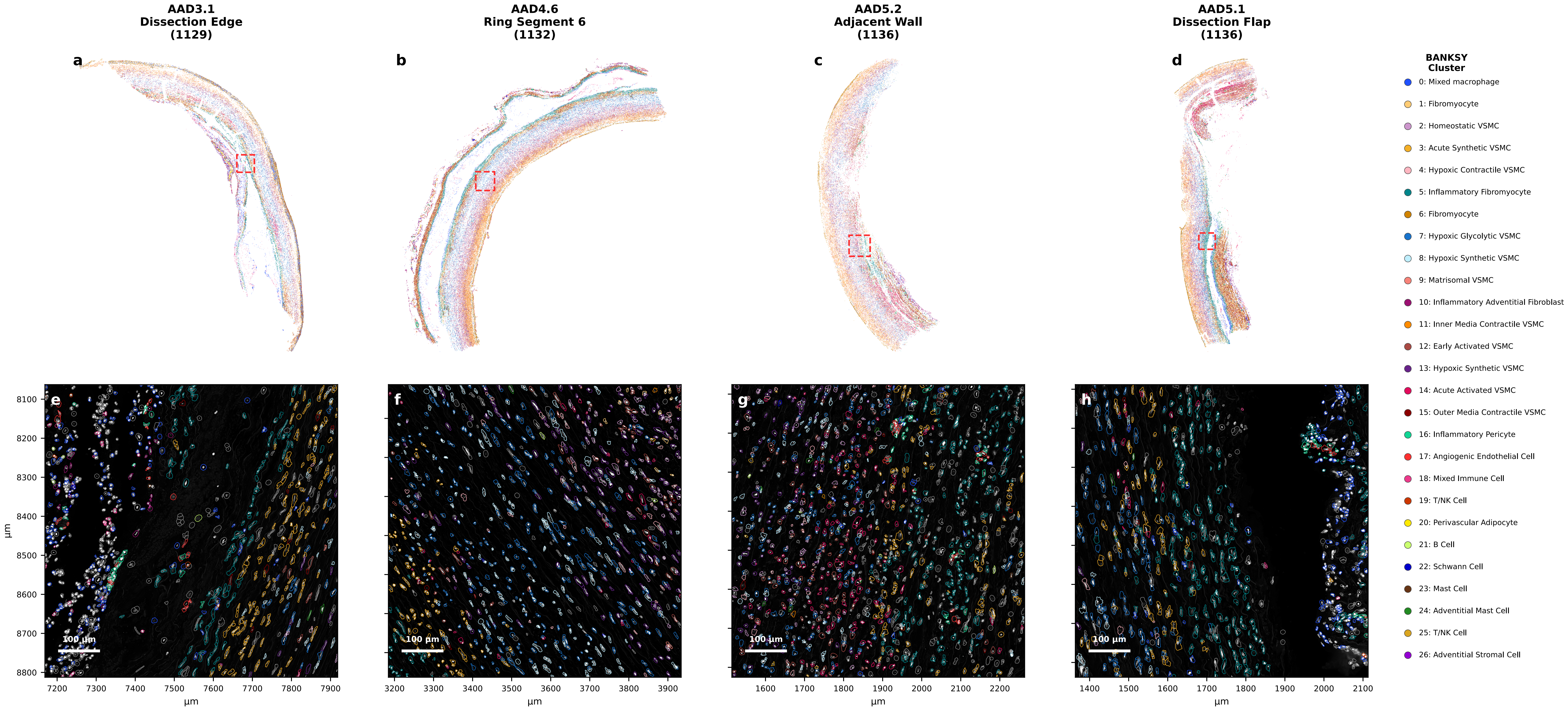


*(a–d) Full-tissue spatial scatter; each dot = one post-QC cell coloured by BANKSY cluster (0–26). Dashed red box = 750 × 750 µm region magnified in (e–h). (e–h) DAPI image with Baysor cell-polygon outlines coloured by BANKSY cluster (see legend), confirming accurate single-cell segmentation morphology across tissue contexts. QC: avg_confidence ≥ 0.5, ≥ 20 counts, ≥ 15 genes, elongation ≤ 99th-percentile (1,215,720 / 1,273,889 cells retained).*

**Figure 4c — Segmentation QC: Full-Tissue Overview and DAPI Zoom Validation**

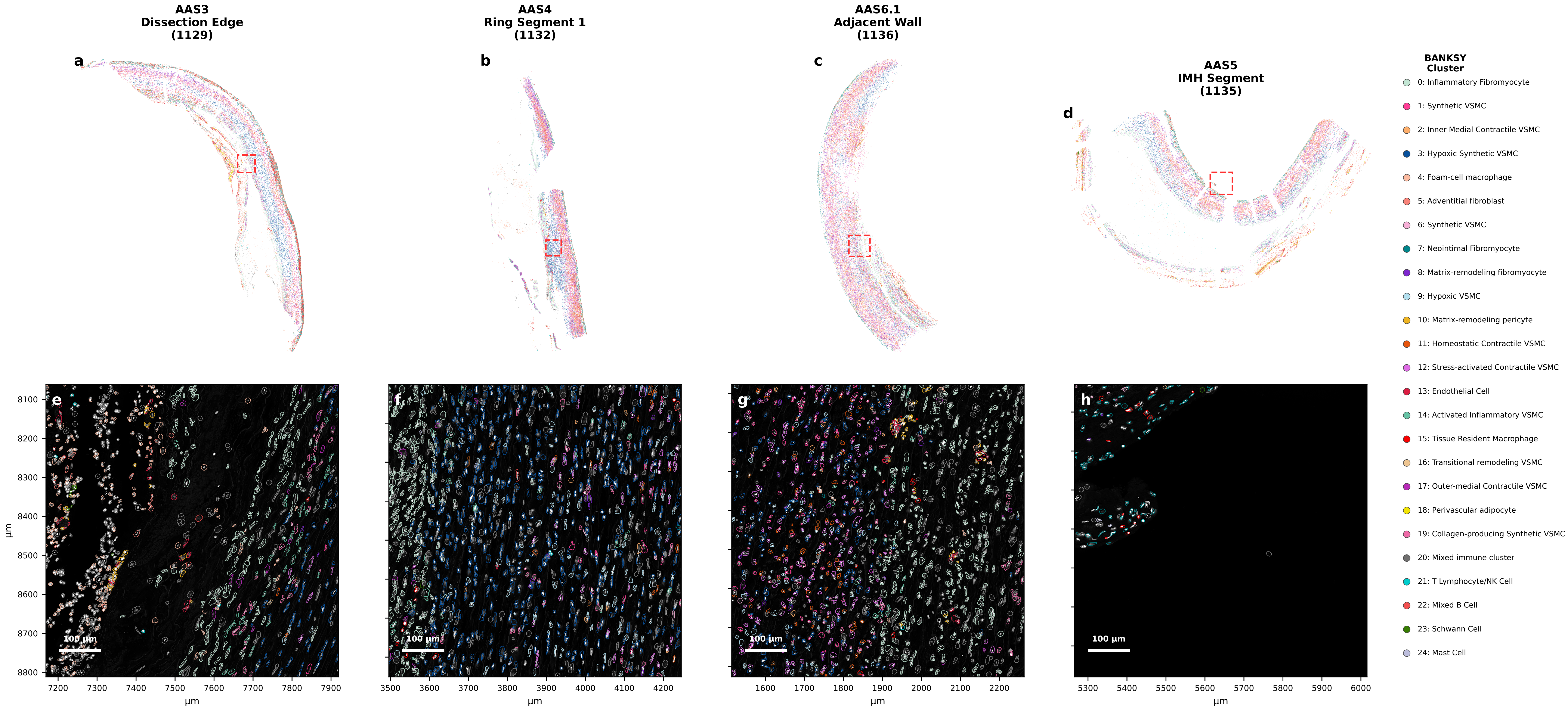


*(a–d) Full-tissue spatial scatter; each dot = one post-QC cell coloured by BANKSY cluster (retained clusters only). Dashed red box = 750 × 750 µm region magnified in (e–h). (e–h) DAPI image with Baysor cell-polygon outlines coloured by BANKSY cluster (see legend), confirming accurate single-cell segmentation morphology across tissue contexts.*

### Supplemental Table — Cell accounting and median wall depth
#### Diseased — 13-sample subset (everything else)

| Sample | Total | Dropped (outside ROI) | Interested (in-domain) | QC-dropped | QC-ok | Median depth (v_equivol) | Median depth (µm) |
|---|---|---|---|---|---|---|---|
| AAS1 | 59,876 | 41,525 | 18,351 | 7,477 | 10,874 | 0.320 | 440 |
| AAS1.1 | 107,319 | 70,154 | 37,165 | 11,856 | 25,309 | 0.463 | 630 |
| AAS2 | 100,486 | 86,348 | 14,138 | 5,275 | 8,863 | 0.539 | 480 |
| AAS3 | 97,475 | 67,104 | 30,371 | 12,874 | 17,497 | 0.390 | 390 |
| AAS3.1 | 183,330 | 145,641 | 37,689 | 13,153 | 24,536 | 0.432 | 950 |
| AAS4 | 84,581 | 43,073 | 41,508 | 16,972 | 24,536 | 0.456 | 660 |
| AAS4.1 | 102,945 | 67,849 | 35,096 | 14,426 | 20,670 | 0.428 | 420 |
| AAS5 | 86,843 | 39,776 | 47,067 | 12,364 | 34,703 | 0.469 | 540 |
| AAS6 | 140,289 | 98,624 | 41,665 | 15,453 | 26,212 | 0.363 | 505 |
| AAS6.1 | 101,056 | 56,202 | 44,854 | 15,201 | 29,653 | 0.428 | 525 |
| AAS7 | 89,955 | 70,383 | 19,572 | 6,277 | 13,295 | 0.519 | 650 |
| AAS8 | 108,511 | 76,295 | 32,216 | 13,017 | 19,199 | 0.439 | 625 |
| AAS8.1 | 105,514 | 74,581 | 30,933 | 10,980 | 19,953 | 0.365 | 500 |
| **Total / pooled median** | **1,368,180** | **937,555** | **430,625** | **155,325** | **275,300** | **0.428** | **540** |

**Supplemental Table – QC Metrics Summary (15 samples, res=1.2)**
**(before QC = raw segmentation output; after QC = postqc_final.h5ad)**

| Sample | Cells (pre-QC) | Cells (post-QC) | Retained (%) | Med. counts (pre) | Med. counts (post) | Med. genes (post) |
|---|---|---|---|---|---|---|
| AAS1 | 59,876 | 28,617 | 47.8% | 19 | 37 | 26 |
| AAS1.1 | 107,319 | 53,088 | 49.5% | 20 | 38 | 26 |
| AAS2 | 100,486 | 62,558 | 62.3% | 31 | 56 | 33 |
| AAS2.1 | 66,588 | 37,122 | 55.8% | 25 | 45 | 29 |
| AAS3 | 97,475 | 58,255 | 59.8% | 31 | 61 | 36 |
| AAS3.1 | 183,330 | 121,316 | 66.2% | 48 | 76 | 40 |
| AAS4 | 84,581 | 45,936 | 54.3% | 22 | 41 | 27 |
| AAS4.1 | 102,945 | 62,369 | 60.6% | 27 | 53 | 33 |
| AAS5 | 86,843 | 52,424 | 60.4% | 27 | 50 | 31 |
| AAS6 | 140,289 | 85,739 | 61.1% | 27 | 50 | 31 |
| AAS6.1 | 101,056 | 58,623 | 58.0% | 24 | 47 | 30 |
| AAS7 | 89,955 | 54,517 | 60.6% | 32 | 58 | 36 |
| AAS7.1 | 61,924 | 38,360 | 62.0% | 44 | 80 | 43 |
| AAS8 | 108,511 | 62,722 | 57.8% | 26 | 47 | 30 |
| AAS8.1 | 105,514 | 59,149 | 56.1% | 24 | 46 | 28 |

#### Supplemental Table — Cell accounting and median wall depth
#### Donor — 5 samples

| Sample | Total | Dropped (outside ROI) | Interested (in-domain) | QC-dropped | QC-ok | Median depth (v_equivol) | Median depth (µm) |
|---|---|---|---|---|---|---|---|
| DC1 | 78,135 | 52,858 | 25,277 | 9,579 | 15,698 | 0.486 | 870 |
| DC2 | 83,513 | 48,399 | 35,114 | 12,743 | 22,371 | 0.419 | 735 |
| DC3 | 57,103 | 39,845 | 17,258 | 8,663 | 8,595 | 0.310 | 400 |
| DC4 | 128,326 | 84,806 | 43,520 | 9,861 | 33,659 | 0.549 | 1,245 |
| DC5 | 69,024 | 18,475 | 50,549 | 10,444 | 40,105 | 0.357 | 610 |
| **Total / pooled median** | **416,101** | **244,383** | **171,718** | **51,290** | **120,428** | **0.438** | **790** |

**Table — QC Filtering Summary (15 samples, res=1.2)**
**Thresholds: counts ≥ 20, genes ≥ 15 | Total pre-QC: 927,176 → post-QC: 888,992**

| Sample | Cells pre-QC | Cells post-QC | Removed | Retained (%) | Med counts (post) | Med genes (post) | Med area (µm²) | Med conf. |
|---|---|---|---|---|---|---|---|---|
| AAS1 | 29153 | 27372 | 1781 | 93.9 | 39.0 | 27.0 | 1435.0 | 0.991 |
| AAS1.1 | 54660 | 51131 | 3529 | 93.5 | 40.0 | 27.0 | 1501.0 | 0.987 |
| AAS2 | 65565 | 63159 | 2406 | 96.3 | 56.0 | 33.0 | 876.6 | 0.977 |
| AAS2.1 | 40410 | 37133 | 3277 | 91.9 | 46.0 | 29.0 | 1286.0 | 0.963 |
| AAS3 | 63963 | 62182 | 1781 | 97.2 | 56.0 | 35.0 | 994.5 | 0.952 |
| AAS3.1 | 128806 | 125174 | 3632 | 97.2 | 75.0 | 39.0 | 888.8 | 0.964 |
| AAS4 | 46224 | 43723 | 2501 | 94.6 | 43.0 | 28.0 | 1485.0 | 0.996 |
| AAS4.1 | 63075 | 60826 | 2249 | 96.4 | 55.0 | 34.0 | 1032.0 | 0.992 |
| AAS5 | 54234 | 52060 | 2174 | 96.0 | 51.0 | 31.0 | 1095.0 | 0.985 |
| AAS6 | 87491 | 83808 | 3683 | 95.8 | 51.0 | 32.0 | 1237.0 | 0.988 |
| AAS6.1 | 59121 | 55555 | 3566 | 94.0 | 49.0 | 31.0 | 1088.0 | 0.992 |
| AAS7 | 60900 | 59487 | 1413 | 97.7 | 53.0 | 34.0 | 1188.0 | 0.938 |
| AAS7.1 | 43553 | 42324 | 1229 | 97.2 | 70.0 | 39.0 | 811.5 | 0.924 |
| AAS8 | 67590 | 65428 | 2162 | 96.8 | 45.0 | 29.0 | 1178.0 | 0.948 |
| AAS8.1 | 62431 | 59630 | 2801 | 95.5 | 46.0 | 28.0 | 1419.0 | 0.973 |
