## Supplementary figures and images for "Cellular-Resolution Spatial Transcriptomics Reveals Laminar VSMC Phenotypic Remodeling and a Hypoxic Medial Core in Human Thoracic Aortic Dissection"

### Supplemental Figures

#### Supplemental Figure 1: Uniform Manifold Projection (UMAP) for AAS samples with all clusters

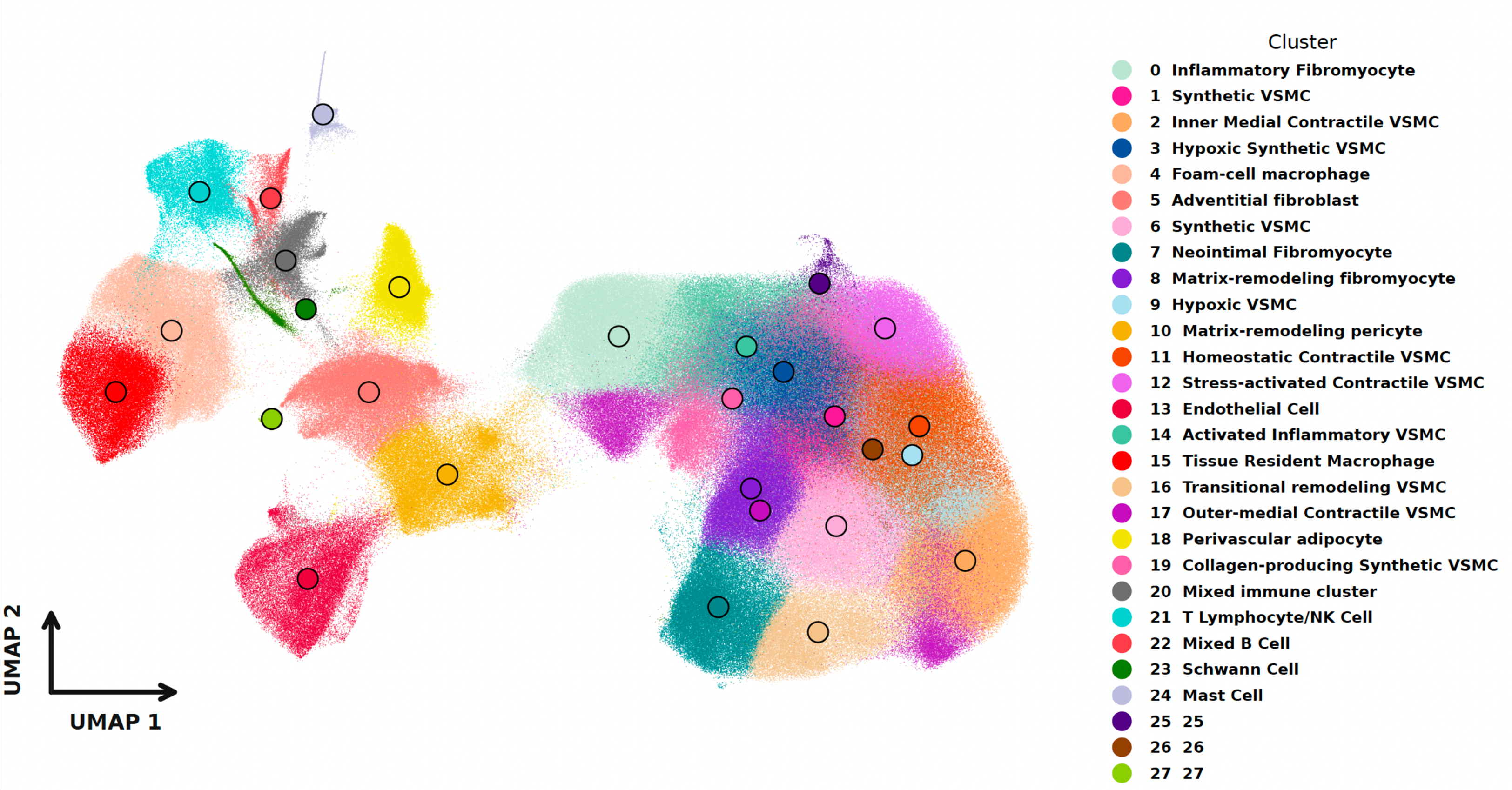
