## Supplemental Tables for "Cellular-Resolution Spatial Transcriptomics Reveals Laminar VSMC Phenotypic Remodeling and a Hypoxic Medial Core in Human Thoracic Aortic Dissection"

**Supplemental Method**

Add a brief paragraph defining the integration metrics: "Integration quality was assessed using batch silhouette score, iLISI, and cLISI; higher iLISI indicates better batch mixing and lower cLISI indicates higher cluster purity.

**Supplemental Table *** — Computational parameter**

| Step | Parameter | Value |
| --- | --- | --- |
| Transcript filtering | Q‑score threshold | ≥ 20 |
| Segmentation | Tool (version) | Baysor v0.6.2 |
| Segmentation | Nucleus boundaries | Xenium default segmentation |
| Segmentation | Scale | 20 px / 4.25 μm |
| Segmentation | Confidence | 0.8 |
| Segmentation | Min transcripts/cell | 30 |
| Cell QC | Total counts | ≥ 20 and ≤ 450 |
| Cell QC | Min genes/cell | ≥ 10 |
| Normalization | Target sum | 100 |
| Normalization | Transform | log1p |
| Batch correction | Tool | Harmony (harmonypy v0.2.0) on PC embedding |
| Integration QC | Metrics | batch silhouette, iLISI, cLISI |
| Spatial embedding | Tool | BANKSY (pyBanksy v1.3.4) |
| Spatial embedding | Neighbors (k) | 15 |
| Spatial embedding | Weight decay | scaled Gaussian, λ = 0.2 |
| Dimensionality | PC selection | Perpendicular distance elbow |
| Dimensionality | PCs retained | 20 |
| Clustering | Method | Leiden on Harmony‑corrected BANKSY embedding |
| UMAP | Neighbors (k) | 50 |
| UMAP | Metric / backend | Euclidean / pynndescent |
| Dataset | Total cells | ~1.2 million |
| Trajectory | Connectivity | PAGA, threshold ≥ 0.20 |
| Trajectory | Diffusion map | n = 15 |
| Trajectory | DPT components | top 10 |
| Trajectory | Root | cell nearest contractile VSMC centroid |
